# Disentangling the lipid divide: Identification of key enzymes for the biosynthesis of unusual Membrane-spanning and Ether lipids in Bacteria

**DOI:** 10.1101/2022.04.21.487849

**Authors:** Diana X. Sahonero-Canavesi, Melvin Siliakus, Alejandro Abdala Asbun, Michel Koenen, F. A. Bastiaan von Meijenfeldt, Sjef Boeren, Nicole J. Bale, Julia C. Engelman, Kerstin Fiege, Lora Strack van Schijndel, Jaap S. Sinninghe Damsté, Laura Villanueva

## Abstract

Bacterial membranes are composed of fatty acids (FAs) ester-linked to glycerol-3-phosphate, while archaea possess membranes made of isoprenoid chains ether-linked to glycerol-1-phosphate. Many archaeal species organize their membrane as a monolayer of membrane-spanning lipids (MSLs). Exceptions to this ‘lipid divide’ are the production by some bacterial species of (ether-bound) MSLs, formed by tail-tail condensation of fatty acids resulting in the formation of (*iso*) diabolic acids (DAs), which are the likely precursors of paleoclimatological relevant branched glycerol dialkyl glycerol tetraether molecules. However, the enzymes responsible for their production are unknown. Here, we report the discovery of bacterial enzymes responsible for the condensation reaction of fatty acids and for ether bond formation, and confirm that the building blocks of *iso*-DA are branched *iso*-FAs. Phylogenomic analyses of the key biosynthetic genes reveal a much wider diversity of potential MSL (ether)-producing bacteria than previously thought, with significant implications for our understanding of the evolution of lipid membranes.

---

Cells are separated from the surrounding environment by a cytoplasmic membrane composed of lipids and proteins. The typical bacterial lipid membrane consists of fatty acids bound to a glycerol-3-phosphate backbone (G3P) via ester linkages, organized in a bilayer structure. Strikingly, however, some bacterial groups organize their membranes in a monolayer of membrane-spanning lipids (MSL) formed by long-chain dicarboxylic acids that are linked to G3P through ester and, sometimes, ether bonds. Both MSL and ether bonds have been considered archetypical archaeal membrane features. Known bacterial MSLs are constituted by diabolic acids (DAs) and *iso*-diabolic acids (*iso*-DAs) (Fig. 1a, Supplementary Fig. 1). DAs have been encountered in members of the phyla Thermotogae and Firmicutes of the Clostridia class^1-4^. *Iso*-DAs occur in members of the genus *Thermoanaerobacter* (phylum Firmicutes, Clostridia class)^5^, as well as in species of the subdivisions (SDs) 1, 3, 4 and 6 of the phylum Acidobacteria^6,7^. DAs have been shown to be biosynthesized by tail-to-tail condensation of two C_16:0_ fatty acids (FAs) at the ω-1 positions^8^. In the same way, *iso*-DAs are believed to be produced by the condensation of two *iso*-C_15:0_ FAs at the ω positions^1^, but the enzymes responsible for the formation of these types of bacterial MSLs (i.e., MSL synthases) remain elusive. A recent study has identified a radical SAM enzyme (tetraether synthase, Tes) responsible for the archaeal tail-to-tail coupling of two ether-bound phytanyl chains, enabling the synthesis of glycerol dialkyl glycerol tetraethers (GDGTs)^9^. Since Tes homologs were detected in bacterial genomes, they have been hypothesized to be involved in the synthesis of bacterial MSLs^9^. However, no Tes homologs were detected in genomes of the Thermotogae known to be MSL-producers^9^, which suggests another enzyme is involved in the synthesis of bacterial MSLs.

**Figure 1.**
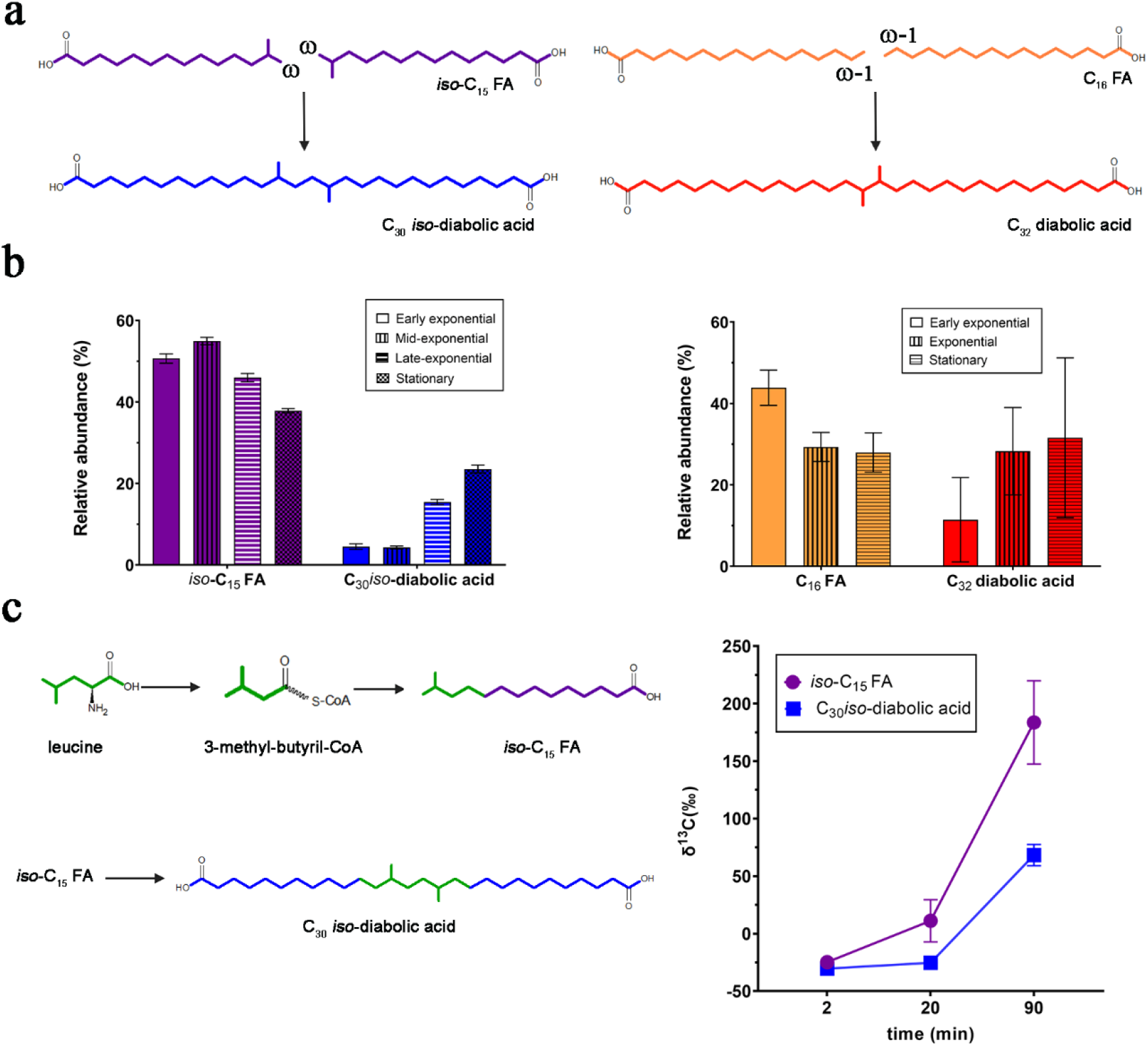
Membrane-spanning lipids are produced via condensation of fatty acids. (a) The C_30_ *iso*-diabolic acid is thought to be formed by coupling two *iso*-C_15_ FAs at their ultimate (ω) carbon atoms. The synthesis of C_32_ diabolic acid proceeds through coupling of two *n*-C_16_ FAs at the penultimate (ω-1) carbon atoms. (b) Relative abundance (%) of core lipids of *iso*-C_15_ and *iso*-diabolic acid C_30_ in cultures of *T. ethanolicu*s grown at 60°C across growth phases (see Supplementary Table 1 and Supplementary Information) and of C_16_ and diabolic acid C_32_ in cultures of *T. maritima* grown at 80°C across growth phases (see Supplementary 2, Supplementary Table 3, and Supplementary Information for further details). In both experiments the relative abundance of MSLs increases with growth. All experiments were performed in triplicate and the error bars indicate ± SD. (c) Labelling experiment using ^13^C-labeled leucine added to cultures of *T. ethanolicus*, lead to the formation of labeled *iso*-C_15_, and subsequently to labeled C_30_ *iso*-diabolic acid in a time-course experiment (see Supplementary Table 2 and Supplementary Information). The degree of labelling is indicated by their δ^13^C values. The leucine-derived carbon atoms (in green) form a part of the carbon skeleton of the C_30_ *iso*-diabolic acid. All experiments were performed in triplicate and the error bars indicate ± SD.

Ether-bonded membrane lipids are also a typical archaeal feature. Nevertheless, they have been found in some bacteria, sometimes together with MSLs (in Thermotogae and several Acidobacteria SDs). Non-isoprenoid alkyl glycerol ether lipids have been found in (hyper)thermophilic species of the bacterial phylum Thermotogae^4^ in aerobic and facultative anaerobic mesophilic bacteria of the Acidobacteria SD1 and 4^6,7^, in *Aquifex pyrophilus*^10^, in *Ammonifex degensii* (Firmicutes Clostridia^11^), in some *Planctomycetes*^12^ and in some sulfate-reducing bacteria^13,14^. In addition, alkenyl (1-alk-1′-enyl, vinyl) glycerol ether lipids or so-called plasmalogens have been detected in non-thermophilic bacteria and suggested to play a role in cell resistance against environmental stresses^15-16^. Enzymes involved in bacterial ether lipid biosynthesis have been discovered in select taxa. In myxobacteria, two independent pathways contributing to the biosynthesis of ether lipids have been identified; the gene product of Mxan_1676 coding for an alkylglycerone-phosphate synthase (*agps* gene) and the *elbB*-*elbE* gene cluster^17^, which has also been detected in SD4 Acidobacteria^7^. A gene encoding a plasmalogen synthase (*pls*A) has been identified in anaerobic bacteria^18^. A modified form of *pls*A has been detected in *Thermotoga maritima* and in other bacteria producing ether-derived lipids^19^ and proposed to be involved in the conversion of bacterial ester bonds into ether bonds generating saturated alkyl ethers.

The reason why bacteria synthesize membrane-spanning and ether lipids, and how these features were acquired, remains poorly understood, but it has been speculated that both the presence of ether bonds and a membrane organization based on a monolayer of MSLs confer membrane stability^20,21^, as shown for archaeal GDGTs^22^. Determination of how bacterial ether lipids and MSLs are synthesized is important to better understand how the divergence of lipid membranes, or the ‘lipid divide’, proceeded in all life forms. In addition, *iso*-DAs are thought to be the main precursors of the branched GDGTs (brGDGTs^23^) (Supplementary Fig. 1), which occur widespread in the environment and are widely used for paleoclimatological reconstructions^24^. However, their biological producers remain unclear. Determining the biosynthetic pathway of bacterial (ether) MSL synthesis will allow for the detection of this capacity in other microbial groups.

Here, we identified and confirmed the activity of an MSL synthase in bacteria. In addition, we confirm the enzymatic activity of a plasmalogen synthase homolog that is involved in the formation of ether bonds in bacterial alkyl glycerol lipids. Based on phylogenetic analyses of these enzymes, we identified microbial groups that have the potential to generate these membrane components and how this feature was acquired, with evolutionary implications for understanding the acquisition of membrane lipids in all life forms.

## *Iso*-diabolic acid is produced via condensation of *iso*-fatty acids

The biosynthesis of the MSL *iso*-DA is thought to proceed through the coupling of the tails of *iso*-branched FAs precursors^1^ (Fig. 1a). Since growth temperature affects membrane stability, and therefore changes in the proportion of MSL are expected, we performed culturing experiments with the *iso*-DA producer *Thermoanaerobacter ethanolicus* under different growth regimes and measured its lipids. The relative abundance of the dominant *iso-*C_15:0_ FA decreased simultaneously with a marked increase in the C_30_ *iso*-DA during growth at optimal 60°C temperature (Fig. 1b). A similar observation was made at suboptimal (45°C) temperature, albeit the maximum abundance of the C_30_ *iso*-DA remained lower (14 *vs*. 24% of total core lipids; Supplementary Table 1). The higher relative abundance of *iso*-C_30_ DA during the stationary phase at optimal *vs*. suboptimal growth temperature suggests that *iso*-DA production is regulated by temperature. This is in agreement with an increased production of GDGTs in archaea at higher temperatures^25^. These results moreover strongly suggested the formation of the C_30_ *iso*-DA proceeds through a coupling of two *iso-*C_15:0_ FAs. To confirm this substrate-product relationship, we performed incubations with *T. ethanolicus* with a labeled branched amino acid, ^13^C-leucine, required for the synthesis of 3-methylbutyryl-CoA, a key building block of *iso*-C_15:0_ FA^26^. Label incorporation was detected in *iso-*C_15:0_ FA but not in C_30:0_ *iso-* DA 20 min after the addition of ^13^C-leucine, whereas after prolonged incubation (90 min) the label was also incorporated into the C_30_ *iso*-DA (Fig. 1c, Supplementary Table 2). These results confirm that *iso-*C_15:0_ FA acts as the precursor for C_30_ *iso*-DA.

## In search for potential proteins for the biosynthesis of membrane-spanning and ether lipids in bacteria

Our results with *T. ethanolicus* indicate that the biosynthetic reaction leading to *iso*-DA is growth phase-dependent (Fig. 1, Supplementary Figure 2, Supplementary Table 3), as recently also shown for the formation of the MSL DA in *T. maritima*^19^ (Fig. 1b, supplementary Table 3, REF^19^). In both *T. ethanolicus* and *T. maritima*, the percentage of MSLs substantially increased during the stationary phase of growth, suggesting that experimental conditions allow for the detection of the activation of the genes coding for the proteins involved in the synthesis of MSLs. To test this, we analyzed the transcriptomic and proteomic response of these two bacterial species and compared them between different growth phases and at optimal and suboptimal growth temperatures (Supplementary Tables 4-14, Supplementary Figures 3-8. Supplementary Information). Supporting the usability of this approach, the gene encoding the modified-*plsA* (Tmari_0479), suspected to be an alkyl ether lipid synthase based on protein homology^19^, was indeed found to be upregulated at a variety of conditions, coinciding with higher proportion of alkyl ether lipids (Supplementary Table 14, Supplementary Figure 8). This is fully in line with its presumed role in the production of ether lipids.

We also searched for potential genes encoding MSL synthases. The anticipated biochemical mechanism for MSL synthesis is based on the dimerization of the FA building blocks through the formation of a carbon-carbon bond between either the ω -1 carbon of C_16_ FAs or the ω carbon of *iso*-C_15:0_ FAs to form DA or *iso*-DA, respectively^8,1^. A radical reaction mechanism for the formation of MSLs in bacteria has been previously proposed^27^, and recently confirmed for the synthesis of the isoprenoid MSLs in Archaea^9^. This reaction would involve a radical intermediate formed by a hydrogen extraction at the tail of one of the fatty acids, followed by a condensation with the other tail, involving the loss of another hydrogen, resulting in the formation of a C-C bond in the absence of an activated intermediate. Such unique reactions are commonly catalyzed by radical proteins, a group that shares an unusual Fe-S cluster associated with the generation of a free radical by the reduction of S-adenosylmethionine (SAM)^28^. Based on these considerations, we defined selection criteria for the detection of potential MSL synthases in the pool of genes found to be activated either in the *T. maritima* transcriptome or proteome, or in the *T. ethanolicus* transcriptome, but not attributed to any known metabolic pathway. These criteria were that the gene or protein (1) should code for a radical-SAM protein that contained the cysteine-rich motif; (CxxxCxxC) normally found in the active site of radical enzymes (2) should encode for an oxidoreductase utilizing an [Fe4 4S] cluster, which can act on CH or CH_2_ groups, (3) should encode a membrane-bound protein, as most of the proteins known to be involved in the formation of core membrane lipids are membrane-associated, and (4) should be homologous to proteins in other MSL producing bacteria (Supplementary Figure 9, 10, Supplementary Tables 15-19, Supplementary Information for details). This resulted in a list of genes encoding potential MSL synthases (Supplementary Tables 16, 17). Additionally, we performed homology searches (protein blast) using the confirmed Tes homolog of the archaeon *Methanococcus aeolicus* (Maeo_0574, ABR56159.1) as query, and a homolog was detected in the genome of *T. ethanolicus* (EGD50779, e-value 1e^-72^ and 24% identity), which was also included in our list of potential bacterial MSL synthases (Supplementary Table 16). Again, no Tes homologs were detected in the genome of *T. maritima*.

## Confirmation of the activity of the potential MSL synthases and ether-lipid forming enzyme

To test the activity of the upregulated radical proteins selected as potential MSL synthases in *T. maritima* and in *T. ethanolicus*, the genes were cloned in an inducible expression vector and expressed in *E. coli* BL21 DE3. This *E. coli* strain produces phosphatidylglycerol (PG) intact polar lipids (IPLs)^29^, thought to be the required building blocks for the MSL synthase of *T. maritima*^19^. This *E. coli* strain does not produce the *iso*-FA building blocks for *iso*-DA but has *n*-C_16:0_, *n*-C_18:1_, and the cyclopropyl FAs cy-C_17:0_ and cy-C_19:0_ as the most abundant FAs. In contrast to earlier suggestions of the involvement of archaeal Tes homologs in the formation of bacterial MSL^9^, the heterologous gene expression of the identified Tes homolog of *T. ethanolicus* did not lead to formation of MSLs under either aerobic or anaerobic conditions. Yet, our heterologous gene expression experiments led to the confirmation of another radical SAM protein homologue found in both *T. ethanolicus* and of *T. maritima* genomes (see Supplementary Table 16-18, 20). This gene (named here *mslS*) encodes an MSL synthase enabling the coupling of two FAs. Expression of the *msl*S of *T. ethanolicus* led to the formation of two dicarboxylic acids that were absent in the control experiment (Fig. 2a). Their formation was only observed when the growth and induction of the expression in *E. coli* was done under anaerobic conditions. They were identified as C_33_ and C_34_ diacids containing one and two cyclopropyl moieties, respectively. Their formation can be envisaged by ω ‐ω coupling of two abundant FAs of the *E. coli* strain: an *n*-C_16:0_ FA with a cy-C_17:0_ FA and two cy-C_17:0_ FA, respectively (Fig. 2c, Supplementary Information, Supplementary Figures 10-11). Other diacids were also formed in lower relative abundance (Supplementary Figures 12).

**Figure 2.**
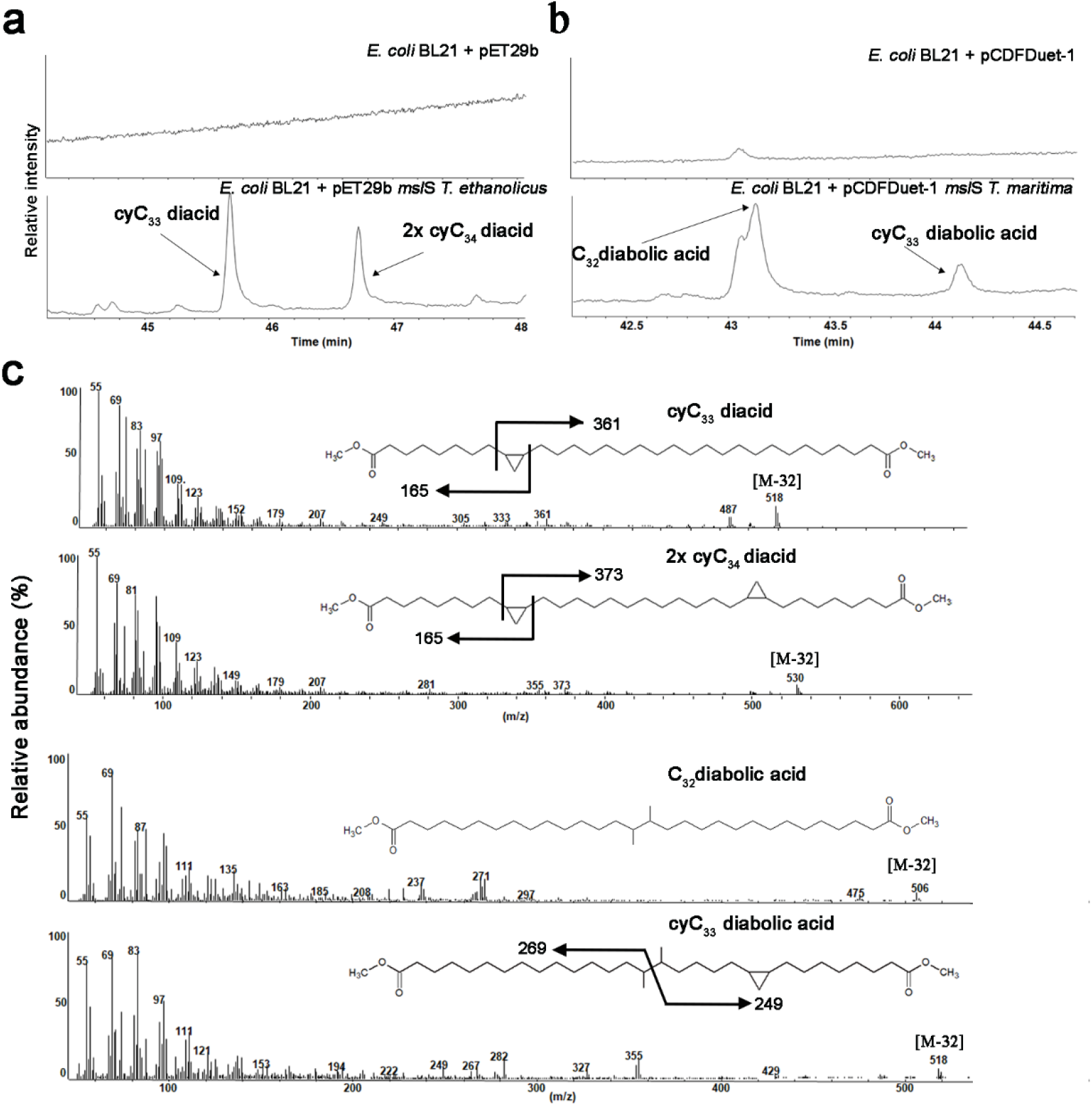
Expression of the bacterial MSL synthases of *T. ethanolicus* and *T. maritima* in *E. coli* results in the production of MSLs through condensation of the tails of two FAs at the ω and ω-1 positions. (a) Partial GC chromatograms (44–48 min) of the base hydrolyzed lipid extract of *E. coli* BL21 (DE3) with ‘empty’ pET29b plasmid (upper trace) or pET29b *msl*S of *T. ethanolicus* plasmid (lower trace), revealing the formation of two major new components, which were identified as C_33_ and C_34_ mono-and bicyclic diacids formed by condensation at the ω-positions of the cyclopropane-containing FAs produced by *E. coli*. (b) Partial GC chromatograms (42-44 min) of the base hydrolyzed lipid extract of *E. coli* BL21 (DE3) with the ‘empty’ pCDFDuet-1 plasmid (upper trace) or pCDFDuet *msl*S of *T. maritima* (lower trace) shows the formation of the C_32_ diabolic acids by C-C bond formation between the ω ‐1 position of two C_16_ fatty acids. (c) Mass spectra of the diacids and diabolic acids formed. To confirm the presence the cyclopropyl moieties the compounds were hydrogenated but remained unaltered.

Similarly, heterologous gene expression of the *msl*S of *T. maritima* in *E. coli* resulted in the formation of C_32_ and C_33_ DAs only under anaerobic conditions (Fig. 2b, Supplementary Figure 13). This result confirms that the expression of *T. maritima mslS* gene product catalyzes the synthesis of C_32_ and C_33_ DA by joining two C_16_ FAs and one C_16_ FA with a cy-C_17:0_ FA respectively at the ω -1 position (Fig. 2c).

Using the same experimental design, we also tested if the expression of the modified *plsA* gene (Tmari_0479 in *T. maritima*) led to the formation of ether lipids in aerobic and anaerobic conditions. Only under anaerobic conditions, the induction of the expression of Tmari_0479 in *E. coli* led to the detection of a series of 1-alkyl glycerol monoethers, where the alkyl chains reflected the major FAs present in the *E. coli* host (Fig. 3, Supplementary Figures 14, 15). 1-Alkyl glycerol monoethers were also detected upon expression in *E. coli* of the modified *plsA* homolog present in the genome of *Desulfatibacillum alkenivorans* (Fig. 3), a non-plasmalogen, ether lipid-producing bacterium^14^ These experiments confirm the earlier proposed function of the modified-plsA in converting an *sn*-1 ester bond into an *sn*-1 ether bond^19^, and we will further refer to this enzyme as glycerol ester reductase (GeR). These findings confirm, for the first time, the enzymatic activity of key enzymes in the production of what are considered to be unusual bacterial membrane-spanning ether lipids.

**Figure 3.**
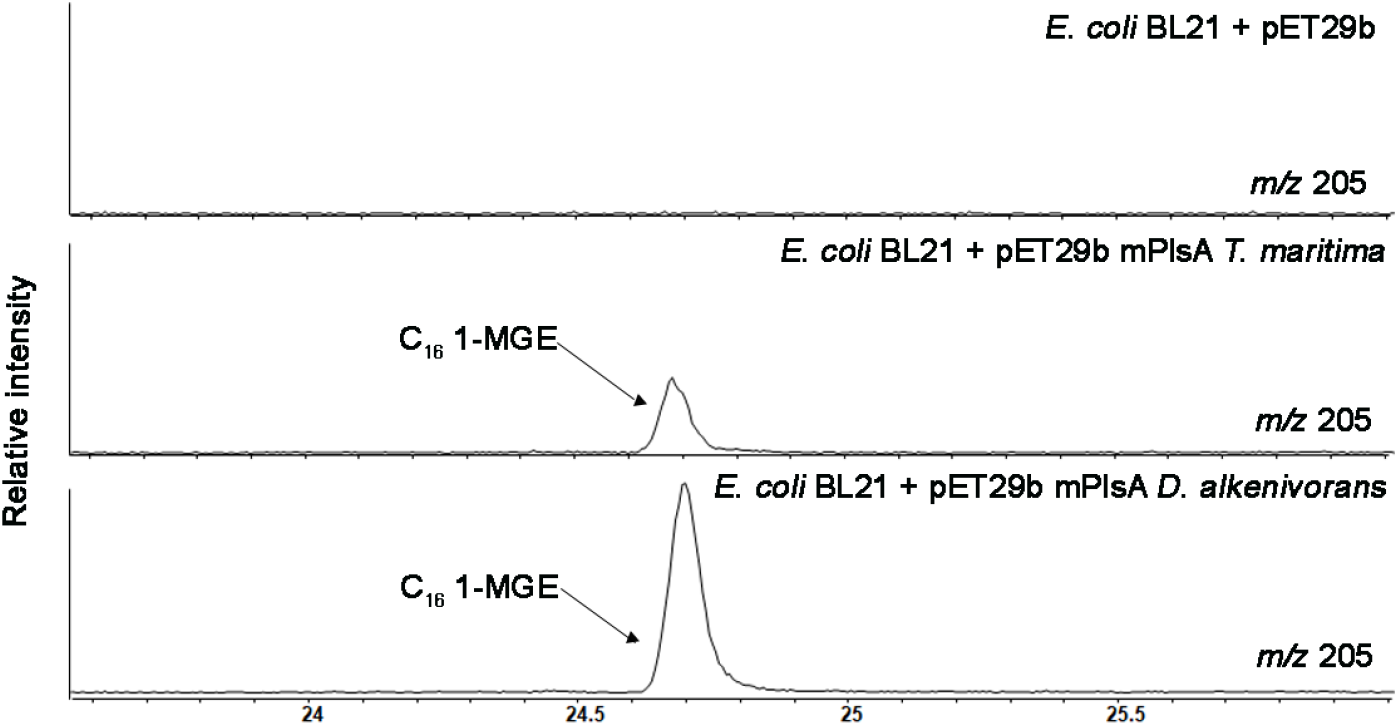
Expression of the modified PlsA of *T. maritima and D. alkenivorans* in *E. coli* results in the production of ether lipids. Partial GC extracted ion chromatograms (*m/z* 205; 24–25.5 min) of the base hydrolyzed lipid extract of *E. coli* BL21 (DE3) with ‘empty’ pET29b plasmid (upper trace) or pET29b containing the mPlsA (Tmari 0479) of *T. maritima* (middle trace) and of *D. alkenivorans* (SHJ90043) (bottom trace) revealing the formation of C_16_ glycerol monoether (MGE, or 1-*O*-hexadecyl glycerol) by *E. coli*

## The widespread occurrence of MSL production in the Domain Bacteria

We screened selected genomes of bacteria with a confirmed presence/absence of DA or *iso*-DA for the presence of homologs of the confirmed MSL synthase. We performed protein Position-Specific Iterative (PSI)-BLAST searches, and we considered as homologs those with an e-value =< 1e^-50^ and identity ≥ 30%. These are stringent search criteria as these proteins belong to the radial SAM family, a large protein family containing many different functions, and less stringent criteria could lead to matches with homologs not involved in the formation of MSLs. The 107 bacterial genomes examined include species of the Clostridia class (phylum Firmicutes), members of the Thermotogales, members of different SDs of the Acidobacteria phylum, as well as others within the Proteobacteria, Chloroflexi, Verrucomicrobia, FCB superphylum, Dictyoglomi, PVC group, Aquificae, and Fusobacteria. These were selected based on their reported membrane lipid composition (presence/absence of MSLs and ether lipids; Supplementary Table 21). These lipid analyses led, for the first time, to the detection of DA biosynthetic capability outside of the Thermotogales and Firmicutes Clostridia, in species within the Dictyoglomi.

Homologs of the MSL synthase were detected in all genomes of the DA-producing Thermotogales and Dictyoglomi phyla, and of the MSL-producing species falling in the Clostridia class (mostly producing *iso*-DAs, except for *Sarcina ventriculi* and *Butyrivibrio fibrisolvens*, both synthesizing DA) (Supplementary Tables 21-22), once again supporting the functionality of the proteins. However, in the Clostridia class the presence of an MSL synthase homolog did not always coincide with confirmed synthesis of MSLs. This suggests that the MSL synthase homolog is not always functional. Alternatively, since we observe that MSL formation is regulated by growth and environmental conditions like temperature, limited experimental setups may have prevented detection of MSLs in these strains. Homologs of Tes were not detected in the genomes of all DA and *iso*-DA producers known to date (Supplementary Table 21), indicating that this protein is not responsible for the biosynthesis of bacterial MSLs in these strains.

Besides, no homologs of the MSL synthase were detected in selected genomes of the Acidobacteria phylum, even though almost all SD 1, 3, 4 and 6 species produce *iso*-DA (Supplementary Table 21). However, this is not unexpected, as most Acidobacteria are aerobic or microaerophilic, and only some species are facultative anaerobes^30^; a strictly anaerobic enzyme such as MSL synthase would therefore likely not be functional. In addition, we only detected Tes homologues in 3 out of the 14 acidobacterial species known produce *iso*-DA, using our stringent search criteria (Supplementary Table 21). Hence, an alternative aerobic pathway used by *iso*-DA producing acidobacteria, which would represent a case of convergent evolution, thus, remains unconfirmed.

An alignment of the MSL synthases of all species with confirmed MSL production with the confirmed MSL synthase of *T. ethanolicus* revealed six conserved blocks and hydrophobic regions near their C-terminus when sequences were grouped according to *iso*-DA (Fig. 4a) and DA producers (Fig. 4b) (Supplementary Figure 16). The novel enzymatic activity leading to the synthesis of MSL either for *iso*-DA or DA consists in joining the alkyl tails of two fatty acids, meaning that the substrates employed by these enzymes are hydrophobic in nature.

**Figure 4.**
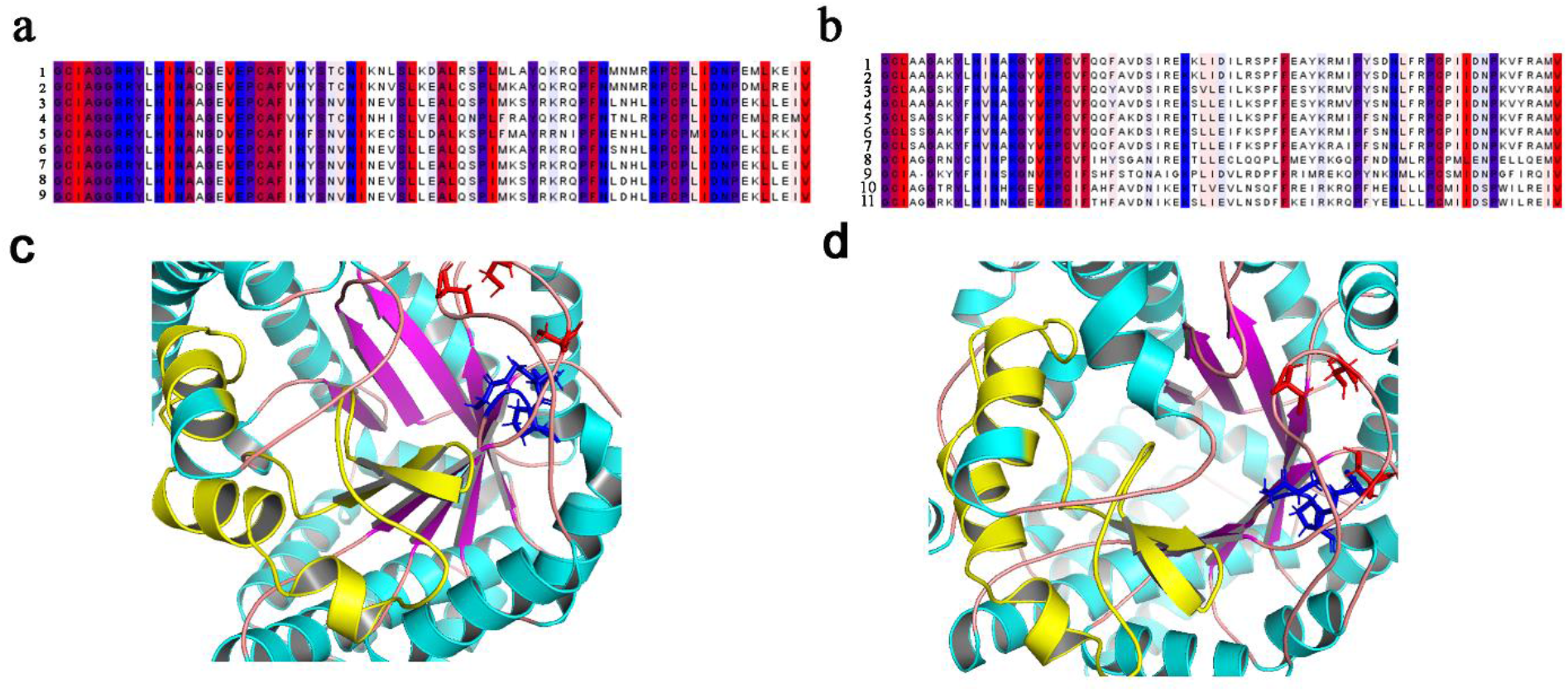
Protein alignment of a section of the two types of bacterial MSL synthases and 3D structures of the MSL synthases from *T. ethanolicus* and *T. maritima*. (a) MSLs producing *iso*-DAs. Key: 1 *Fervidicola ferrireducens*, 2 *Thermosediminibacter oceani*,3 *Thermoanaerobacter wiegelii*, 4*Moorella thermoacetica*, 5 *Caldicellulosiruptor owensensis*, 6 *Caldanaerobacter subterraneus*, 7 *Thermoanaerobacter siderophilus, 8 Thermoanaerobacter thermohydrosulfuricus, 9 Thermoanaerobacter ethanolicus* and (b) MSLs producing DAs. 1 *Thermotoga maritima, 2 Thermotoga neapolitana*, 3 *Pseudothermotoga elfii, 4Pseudothermotoga hypogea, 5 Thermosipho africanus, 6 Thermosipho melanesiensis, 7 Fervidobacterium pennivorans, 8 Butyrivibrio fibrisolvens*, 9 *Sarcina ventriculi*, 10 *Dictyoglomus thermophilum*,11 *Dictyoglomus turgidum*. Predicted amino acid regions colored according to hydrophobicity^72^ the most hydrophobic residues are red and most hydrophilic are colored blue. Lower panel depicts the modelled 3D structures of the MSL synthases of (c) *T. ethanolicus* and of (d) *T. maritima* zoomed in on the region of the conserved cysteines (red), the SAM binding motif (blue) and the region corresponding to the alignment proposed to interact with the lipid substrate (yellow).

Modeling of the 3D-structures of the confirmed MSL synthases of *T. ethanolicus* (Fig. 4c) and *T. maritima* (Fig. 4d) revealed that the proposed hydrophobic region is localized close to the three conserved cysteines and the SAM binding region (radical SAM core), suggesting that this is the reaction center for the coupling of the two hydrophobic tails of the substrates. The key difference between the two types of MSL synthases (i.e., responsible for either DA and *iso*-DA production) is the lipid substrate (i.e., non-branched or *iso*-FA, respectively) and the positions at which the two chains are connected (i.e., ω -1 or ω, respectively). The hydrophobic region of the two groups of MSL synthases likely controls the binding of specific lipid substrates and allows the hydrogen abstraction at the specific position (Fig. 4c,d, Supplementary Figure 17). Indeed, as would be expected for an enzyme forming *iso*-DAs (coupling ω carbon from *iso*-C_15_ FAs), in our expression experiments with the *T. ethanolicus* MSL synthase, several long-chain diacids (Fig. 2a, 2c, Supplementary Figure 8) were produced by the ω ‐ω coupling of two fatty acids, even though only the non-branched FAs produced by the *E. coli* host strain were available for the enzymatic reaction. Consistently, the formation of C_32_ DA in *E. coli* upon expression of the MSL synthase from *T. maritima* confirms the expected enzymatic reaction between the ω - 1 carbons from C_16_ FAs. Moreover, only specific FA combinations resulted in the formation of diacids with *T. ethanolicus* MSL synthase, suggesting that only the tails of specific fatty acids could be accommodated in the hydrophobic region to allow the ω ‐ω coupling. Thus, the *T. ethanolicus* MSL synthase has a well-defined specificity, which is different from the MSL synthases involved in DA synthesis.

We extended the screening of MSL synthase homologs in the NCBI non-redundant protein sequence database (nr) using stringent search criteria (DIAMOND search with e-value <= 1e^-50^ and query coverage ≥ 30%), and built a maximum-likelihood phylogenetic tree. Homologs were detected in species of the phyla Firmicutes, Actinobacteria, Thermotogae, Synergistetes, Spirochaetes, Armatimonadetes, Caldiserica, Calditrichaeota, Coprothermobacterota, Chloroflexi, Nitrospirae, Elusimicrobia, Deferribacteres, Dictyoglomi, Proteobacteria (gamma and delta), Atribacterota, FCB superphylum, Cyanobacteria/Melainabacteria group, PVC group, and in multiple members of the bacteria candidate phyla (see Fig. 5a, Supplementary File 1). The MSL synthase tree clustering does not follow the grouping based on taxonomy, suggesting large scale transfers across the bacterial and archaeal domains. MSL synthase homologs are present in genomes of several classes within the phylum Firmicutes supporting the acquisition of the *msl*S gene before the diversification of this phylum. MSL synthase sequences detected in bacteria candidate phyla, FCB superphylum, Spirochaetes, some Chloroflexi and deltaproteobacteria were closely related, which might suggest they were acquired by horizontal gene transfer as these groups often coexist in anoxic environments.

**Figure 5.**
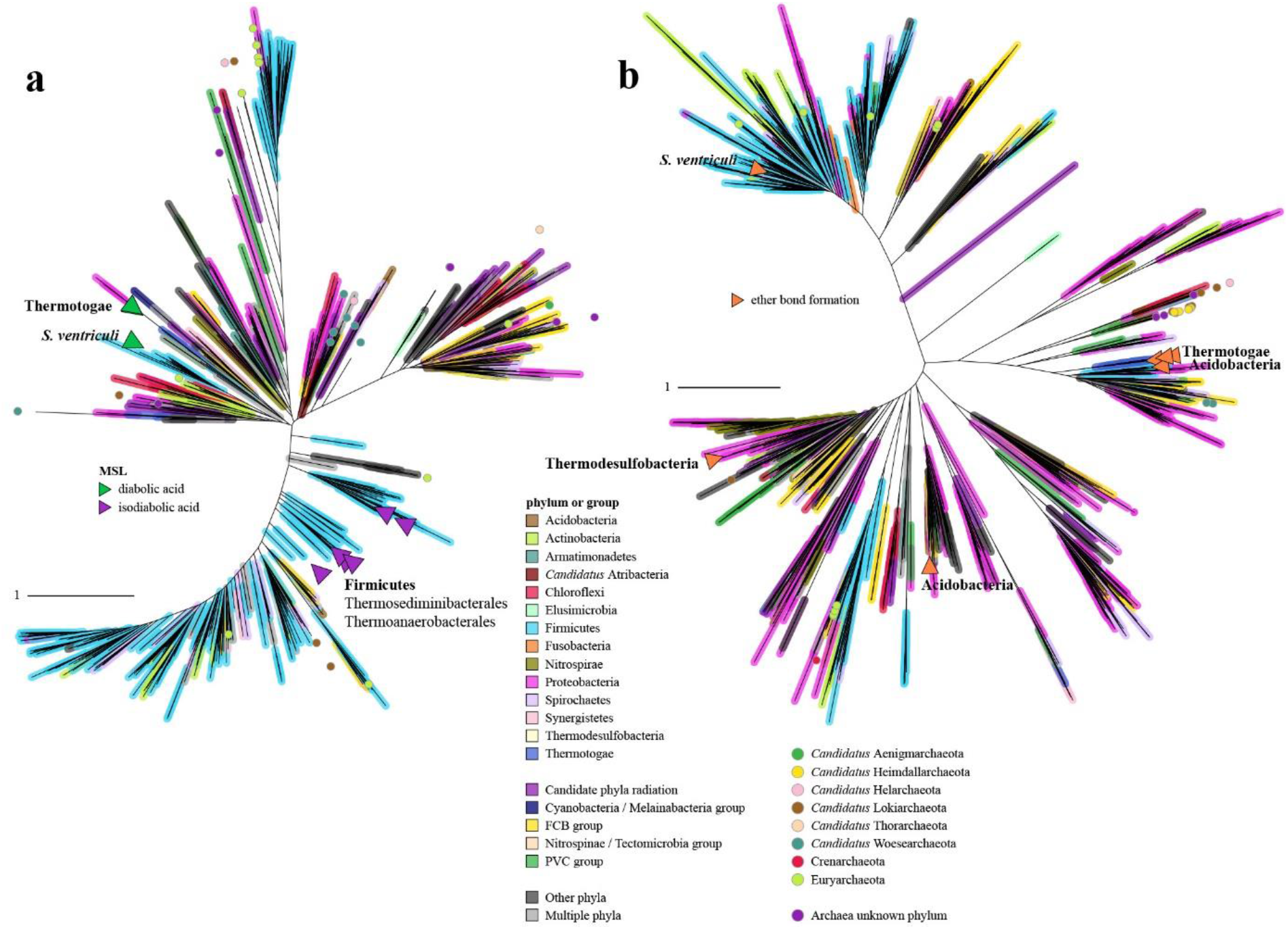
The widespread capability of MSL and ether lipid production in the Bacteria and Archaea domain. Maximum likelihood phylogenetic tree of the predicted homologs of the confirmed (a) MSL synthase and (b) ether bond-forming enzyme, GeR, across the tree of life grouped at the phylum or superphylum level as shown with the colors on the branches and indicated in the legend. Similar sequences were clustered based on sequence identity before tree inference. Purple, green and orange triangles indicate the genomes of microorganisms with homologs that have been seen to synthesize diabolic, *iso*-diabolic or ether-bonds in screened cultures listed in Supplementary Table 21. Colored circles in the tree indicate archaeal taxa. Full trees can be seen in the Supplementary Files 1 and 2. Scale bars represent number of substitutions per site.

## The widespread occurrence of ether lipid production in the Domain Bacteria

In addition to our MSL synthase search, we screened the genomes of a set of selected strains rigorously analyzed for the presence of MSL and ether lipids (Supplementary Table 21), for the presence of homologs of the GeR identified in *T. maritima*. For all of the strains producing alkyl ether-bond membrane lipids, their genomes also harbored a GeR homolog, with the exception of the members of the Acidobacteria. The presence of a GeR homolog does not always lead to confirmed production of alkyl ether lipids in culture (Supplementary Table 21), suggesting that their synthesis may be regulated by specific physiological factors.

The functional domain architecture of the GeR in *T. maritima* is composed of two activation domains identified by the Pfam domain PF01869 (or B; BcrAD_BadFG), one reduction domain PF09989 (or D; DUF2229) followed by four small (*ca*. 50 aa) domains, two reduction (D) and two dehydration domains PF06050 (or H; HGD-D) (BBDHDDH, Supplementary Figure 18). We analyzed the functional domain architecture of the GeR homologs in strains with confirmed production of saturated alkyl ethers (Supplementary Table 21).

The *T. maritima* GeR, confirmed here to form alkyl ethers, coincides in the three first domains with that of *E. faecalis* (BBDH) (Supplementary Figure 18, REF^19^), which is known to synthesize plasmalogens^18^. Other previously screened Thermotogae strains known to synthesize alkyl ethers, harbor a GeR that also coincides with the two activation and reduction domains (BBD), but with some variations in the successive domains (Supplementary Figure 18, Supplementary Table 21). Similarly, the GeR functional domains of *D. alkenivorans*, which synthesizes alkyl ethers, also possess the activation and reduction domains, but also with some variations in the architecture with respect to the functional domain localization of the *T. maritima* GeR.

We observed that several strains within the Clostridia synthesizing plasmalogens, and in lower proportion also saturated alkyl ethers, have the same functional domain architecture present in *Enterococcus faecalis* (BBDH). It is possible that the synthesis of alkyl ethers is a by-product of the reaction leading to plasmalogens in these cases. All these variations detected along the GeR structural domains indicate that the two activation and one reduction domains (BBD) are conserved among the alkyl ether lipid producers as seen in the confirmed GeR enzyme from *T. maritima* and *D. alkenivoran*s. GeR homologs were also found with other domain architectures containing only one activation domain in the protein (B and BDDD; Supplementary Figure 18, Supplementary Table 21). Since most homologs have two activation domains, it is unclear if a single activation domain leads to a functional protein. Bacteria that have a GeR homolog with only one activation domain may then use an alternative pathway for catalyzing ether lipid formation. Here, we also detected distant homologs of the alkylglycerone-phosphate synthase (*agps* gene) Mxan_1676 of *Myxococcus xanthus*, also involved in the biosynthesis of ether lipids^17^ (Lorenzen *et al*., 2014), in some bacterial genomes (Supplementary Table 21), suggesting that in these organisms more than one biosynthetic pathway for ether lipid production may be used. Some members of the Acidobacteria SD4 have been seen to harbor the *elb* gene cluster^7^. Besides, we also detected the GeR homolog in two Acidobacteria genomes of different subdivisions, *Edaphobacter aggregans* (SD1) and in *Holophaga foetida* (SD8), both reported to synthesize alkyl ether bonds (Supplementary Table 21). Of all the Acidobacteria strains known to synthesize ether-bonded lipids, *E. aggregans* and *H. foetida* are either facultative anaerobes or strict anaerobes. We also detected a GeR homolog in the anaerobic acidobacterium *Thermotomaculum hydrothermale*, but it has not been reported to synthesize alkyl ether lipids in culture (Supplementary Table 21) (REF^30^ for an overview). The (facultative) anaerobic metabolism of these acidobacteria would be compatible with the catalysis of GeR, which it is only functional under anaerobic conditions (REF^18^, this study). Therefore, we speculate that facultative or strict anaerobic groups within the Acidobacteria could be producers of ether lipids by using the GeR confirmed in this study. This hypothesis is further supported by the detection of GeR homologs in Acidobacteria metagenomic assembled genomes detected in anoxic systems (see Supplementary Information).

We also investigated the presence of homologs of GeR of *T. maritima* in the NCBI non-redundant protein sequence database (nr) using stringent search criteria (DIAMOND search with e-value <= 1e^-50^ and query coverage ≥ 30%), and they were detected widely across the tree of life, in species of the phyla Firmicutes, Actinobacteria, Thermotogae, Synergistetes, Spirochaetes, Armatimonadetes, Calditrichaeota, Chloroflexi, Nitrospirae, Elusimicrobia, Deferribacteres, Proteobacteria, Atribacterota, FCB superphylum, Cyanobacteria/Melainabacteria group, PVC group, Nitrospinaea, Thermodesulfobacteria, Aquificae, Acidobacteria, Chryosiogenetes, and in multiple members of the bacteria candidate phyla (Fig. 5b, Supplementary File 2). GeR homologs of Firmicutes are widespread across the GeR protein tree but often mostly closely related to those of Actinobacteria. In addition, multiple GeR homologs were detected in genomes of deltaproteobacteria, which suggests that the capacity to make ether-bonded lipids in members of this group is more widespread than originally thought.

Importantly, many taxa contain both MSL synthase and GeR homologs. We conclude that members of the Firmicutes, Actinobacteria, Thermotogae, Synergistetes, Spirochaetes, Armatimonadetes, Calditrichaeota, Chloroflexi, Nitrospirae, Elusimicrobia, Deferribacteres, Proteobacteria, Atribacterota, FCB superphylum, PVC group, and members of several bacteria candidate phyla are potential producers of bacterial ether MSLs, and therefore potential producers of brGDGTs in environmental settings.

## Occurrence of bacterial MSL synthase and GeR in the Domain Archaea

Most Archaea synthesize MSLs based on isoprenoidal alkyl chains (i.e., GDGTs) linked through ether bonds to G1P instead of the G3P backbone observed in Bacteria. Unlike for Bacteria, the enzymes involved in the formation of ether bonds in Archaea are known (i.e., prenyltransferases^31^) and they are not generally found in Bacteria^32^. The enzyme involved in the coupling of two phytanyl chains to form archaeal MSLs (i.e., biphytanes) has been recently confirmed to be a radical SAM protein (the previously mentioned Tes^9^) and hence the biosynthetic pathway for GDGT production in archaea has been established. The presence of homologs of the confirmed ‘bacterial type’ MSL synthase in genomes of Archaea would thus be unexpected. Surprisingly, however, we also detected homologs in (meta)genomes of members of the DPANN Archaea, and members of the Asgard group (Fig. 5a, Supplementary File 1, Supplementary Information). These homologs of the bacterial MSL synthase could potentially be able to connect the tails of two isoprenoid chains since this is reaction is somewhat similar to the one connecting two FAs for the production of bacterial MSLs. However, the genomes of these archaeal species also harbor homologs of Tes^9^. Furthermore, the low homology with the confirmed bacterial MSL synthase, and the apparent limited archaeal distribution of the MSL synthase homologs argues against the canonical coupling of isoprenoids by MSL synthase in archaea. It is possible that the MSL synthase homologs in specific Archaeal groups catalyzes the coupling of ether or ester bound fatty acids, as seen in this study for Bacteria. This capacity is enigmatic in Archaea. In the same way that the formation of MSLs in Bacteria challenges the concept of the ‘lipid divide’, some Archaea have been reported to produce phospholipid fatty acids^33^ (Gattinger *et al*., 2002), or encode fatty acid biosynthesis genes and ester-bond forming acid transferases alongside the archaeal lipid biosynthesis gene machinery in their genomes^34,35^. This trait has been observed in genomes of uncultured archaeal groups, namely the Euryarchaeota Marine Group II and members of the Asgard superphylum^32,35^. As the same members of the Asgard superphylum seem to harbor a bacterial MSL synthase homolog, some members of this superphylum may have the capacity to form bacterial membrane-spanning phospholipid fatty acid lipid membranes. This hypothesis is exciting due to the position of Asgard archaea as the closest descendants of the archaeal ancestor leading to eukaryotes^36^, and its significance for better understanding the acquisition of membranes in the membrane transition in eukaryogenesis. The detection of homologs of the MSL synthase in members of the DPANN archaea is also of interest. Their streamlined genomes very often lack the biosynthetic genes required to synthesize their own membrane lipids, relying on those of their host^37^. Nevertheless, the presence of the homolog of MSL synthase could be interpreted as a potential capacity to remodel the lipids of the host or those potentially acquired from the environment. Further study is required to investigate these hypotheses and their evolutionary implications.

We also encountered homologs of the GeR in genomes of the Euryarchaeota classes Methanosarcinales and Methanomicrobiales, in the genomes of members of the DPANN archaea (*Ca*. Woesearchaeota, Pacearchaeota), and some members of the Asgard group (*Ca*. Lokiarchaeota, including *Candidatus* Prometheoarchaeum syntrophicum^38^, and genomes of the *Ca*. Heimdallarchaeota) (Fig. 5b), for which we have also detected the presence of potential MSL synthase homologs, giving further support to the possibility that specific Archaea might possess the genetic ability to synthesize (ether-bonded) bacterial MSLs.

## Conclusions

We conclude that the capacity of bacterial (ether-based) MSL production is extended to numerous members of bacterial phyla. These findings have wide implications for the biological sources of bacterial brGDGTs (based on *iso*-DA), which are extensively used for paleoclimate reconstruction^24^. Acidobacteria have been thought to be the biological sources of brGDGTs in specific environments (e.g., REF^20^) and have been detected in a few species of Acidobacteria^6,7^. However, several studies have pointed to other, multiple bacterial sources of brGDGTs, especially in anoxic settings where most MSL-producing acidobacteria cannot thrive^39^. In the current study, we predict the synthesis of bacterial ether MSLs by bacterial groups outside of the Acidobacteria, as well as yet-uncultured (facultative) anaerobic acidobacteria, being those commonly present in soils, freshwater and marine systems. This information will be essential to further constrain the interpretations made based on brGDGT distributions. The fact that the enzymes involved in the synthesis of bacterial (ether-based) MSLs are different from those forming GDGTs in Archaea (i.e., Tes enzyme^9^; and archaeal prenyl transferases^31^) indicates that similar membrane lipid features have been acquired independently in the evolution of lipid membranes, but likely emerged due to the same evolutionary pressure (e.g., increase of membrane stability). The potential synthesis of bacterial-type (ether) MSLs by archaeal members of the Asgard group, together with archaeal lipids, could be an adaptation leading to the hypothetical membrane transition from archaeal to bacterial type during eukaryogenesis. Follow-up studies will be needed to determine which physiological/environmental conditions trigger the formation of these membrane lipids and when and how this evolutionary trait was acquired and retained.

## Material and methods

### Strains, media, and growth conditions

*Thermotoga maritima* (strain MSB8, DSMZ-Deutsche Sammlung von Mikroorganismen und Zellkulturen GmbH, DSM 3109) was cultivated in basal media (BM) under anaerobic conditions in 120-ml batch cultures in 250 ml serum bottles. Cultures were incubated either at optimal (80°C) or lower (55°C) growth temperatures, and after three passages, the batch cultures were inoculated from the acclimated bottle. Growth was monitored by measuring the optical density (OD) at 600 nm and by fluorescent microscopy. Samples for proteomic, transcriptomic, or lipidomic analysis were derived from five replicates at each condition, for each analysis. The cells were harvested at early exponential, exponential, and stationary phases by centrifugation at 3500 rpm for 10 min at 4°C. The supernatant was discarded, and the remaining pellet was immediately frozen at - 20°C until further processing for lipid, protein, or RNA extraction.

*Thermoanaerobacter ethanolicus* JW200 (REF^40^) (DMS 2246) was cultivated under anaerobic conditions in 80 ml batch cultures in 250 ml serum bottles. The media was composed of (per liter): (NH_4_)_2_SO_4_ 1.3 g, KH_2_PO_4_ 0.375 g, K_2_HPO_4_ 0.75 g, MgCl_2_ x 6H_2_O 0.4 g, CaCl_2_ x 2H_2_O 0.13 g, FeSO_4_ x 7H_2_O (0.001% (w/v) in H_2_SO_4_ 2 mM), yeast extract 4 g, resazurin standard stock solution 100x, and pH was adjusted to 6.7 with 5 M NaOH. The medium was anaerobically dispensed in 250 ml serum bottles, and a gas phase of N_2_ was applied. After sterilization, individual bottles were supplemented with 20× cellobiose solution (100g/L) and 100× dilution of L-cysteine hydrochloride monohydrate stock. Cultures were grown at 45°C or 60°C for 1 sub-cultivation growth cycle to ensure acclimatization. Growth was monitored by measuring OD_600nm_. Cells were harvested at the early exponential (OD_600nm_ 0.25), mid-exponential (OD_600nm_ 0.4–0.5), late-exponential (OD_600nm_ 0.65–0.73), and stationary (OD_600m_ 0.62–0.67) growth phases. The experiments were performed in triplicate. At given OD_600nm_, cultures of 80 ml were split in two and immediately centrifuged 15 min at 4600x*g* at 4°C. Half of the cultures were used for RNA sequencing, the other half for lipid extractions.

For the time-course ^13^C labelling experiments with *T. ethanolicus* with labeled leucine, cells were anaerobically grown in 40 ml of media with either L-leucine or ^13^C_6_-labeled leucine (Merck Sigma-Aldrich), added to a final concentration of 0.191 mM in the growth media, and samples were collected after 2, 20, and 90 min of incubation at 55°C and harvested according to the above-described procedure.

Strains of the phylum Firmicutes, Clostridia class analyzed for their lipid composition were cultured by DSMZ with their preferred media to stationary phase. Cultures were harvested by centrifugation and cell pellets freeze-dried before lipid analysis.

### Lipid extraction and analysis

Fatty acids and membrane-spanning lipids (DA and *iso*-DA) were extracted from freeze-dried pellets and analyzed as previously described^19^. Core lipids were identified based on literature data and library mass spectra. The double bond positions present in the identified diacids were determined by derivatization with dimethyl disulfide (DMDS)^41^ with some modifications (incubation at 40°C and then overnight, addition of 400 µL *n*-hexane and 200 µL 5% solution Na_2_S_2_O_3_, aqueous layer extracted two times with *n*-hexane), prior to mass spectrometric identification by GC-MS as described by REF^19^.

For the time-course ^13^C labelling experiments with leucine, harvested cells of *T. ethanolicus* were frozen and freeze-dried. Lyophilized cells were hydrolyzed with 1.5N HCl in methanol by refluxing for 3 h and the pH was adjusted to neutral by KOH addition. The FAs and *iso*-DA in the obtained extracts were derivatized to their methyl esters using BF_3_-methanol solution and analyzed by GC-irmMS as previously described^42,43^.

### RNA extraction and transcriptomic analysis

*T. maritima* cell pellets were defrosted on ice and washed with 500 µL of DNase/RNase free water (New England Biolabs). Cells were resuspended in 700 µL of RLT buffer, and 50 mg of acid-washed glass beads (0.1 µm diameter) were added to a safe-lock tube containing the cell suspension. Cell disruption was performed with the OMNI bead mill Homogenizer (6.3 m/s) 2x. After centrifugation and separation of the cell lysate, the RNA was extracted with the RNeasy® Mini Kit (QIAGEN). The RNA concentration was determined using the Qubit® HS protocol (ThermoFisher). 18 µl (2 µg) of total RNA was hybridized with the Pan-Prokaryote riboPOOL 3’-biotinylated probe (siTOOLs Biotech). The rRNA was subsequently depleted with Hydrophilic streptavidin magnetic beads (New England Biolabs). The depleted rRNA sample was purified before library preparation with RNA clean and concentration (Zymo research) and eluted in 15 µl of DNase-/RNase-Free Water. Samples were QC analyzed with the Agilent RNA 6000 Nano Chips and the Agilent 2100 Bioanalyzer system and stored at -80°C until library preparation.

For *T. ethanolicus*, cells cleared from medium were snap-frozen in dry-ice and stored at -80°C until usage. For RNA extraction the frozen cell pellets were resuspended in 4 ml RNA-later (Invitrogen) while maintained on ice. A 200 µl sample was taken from the cell suspension for RNA isolation. The cells were incubated at room temperature for 5 min. The cells were then pelleted for 10 min at 4700x*g*. The cell pellet was resuspended in 100 µl TE buffer pH 7.5. 500 µl TRIzol reagent was added and mixed. The subsequent suspension was transferred to a pre-cooled 2 ml screw-cap tube supplemented with 250 µl sterile 0.1 mm and 1 mm glass beads. Cells were lysed by beating 6 rounds of 25 sec pulse at 6500 rpm and 5 min pause on watery ice. Ice-cold chloroform (200 µl) was subsequently added and mixed by shaking. Two phases were produced by centrifugation for 15 min. at 12,000x*g*. The transparent aqueous phase was transferred to a new RNAse-free pre-cooled tube without disturbing the interphase. Ethanol was added at 1 volume and mixed. The isolate was transferred to an RNAeasy column and centrifuged 15 seconds at 8000 x*g* to bind nucleic acids to the column. The column was rinsed with 350 µl RLT buffer by centrifugation. An 80 µl RNAse-free DNAse reaction mixture was added to the column and incubated 20 min at 30°C to digest DNA contaminants. The column was rinsed with 350 µl RW1, incubated 5 min and centrifuged. The column was then washed with 700 µl RPE, centrifuged and washed with 80% ethanol. The column was dried by a 2 min spin at 21.000 x*g*. To elute the RNA, 35 µl nuclease-free water was added and eluted by 1 min centrifugation at 8,000 x*g*. For rRNA depletion, 18 µl (1 µg) of total RNA was hybridized with the Pan-Prokaryote riboPOOL 3’ biotinylated probe following the depletion protocol and stored at -80°C until library preparation.

For *T. maritima*, we sequenced 30 RNA samples, five biological replicates across three growth phases (early exponential, exponential and stationary) and two temperature conditions (55°C and 80°C). For *T. ethanolicus* we sequenced 18 RNA samples, three biological replicates across three growth phases (mid exponential, late exponential and stationary) and two temperature conditions (45°C and 60°C). rRNA depleted RNA samples were used to prepare sequencing libraries with the TruSeq RNA stranded kit and sequenced at the Utrecht Sequencing facility (USeq, The Netherlands) on an Illumina NextSeq500 sequencing platform in single-end mode with a read length of 75 nt.

Both transcriptomics libraries were treated equally unless specified otherwise. FASTQ files containing the Illumina reads were quality filtered and standard Illumina adapters removed using Trimmomatic v0.36 (REF^44^) (parameter settings: IlluminaClip: TruSeq3-SE.fa:2:30:10, leading: 20, trailing: 20, sliding window: 5: 20 min length: 40); poly-G tails were removed using cutadapt v1.16 with the --nextseq-trim paremeter set to 20 (REF^45^). Gene counts were calculated using Salmon v1.1.0 (REF^46^) for the *T. maritima* MSB8 library and Salmon v1.3 for the *T. ethanolicus* JW 200, both with the mapping-based mode against the reference genome, being *T. maritima* MSB8 (REF^47^) (NCBI Reference Sequence: NC_021214.1) and *T. ethanolicus* JW 200 (NCBI Reference Sequence: CP033580.1) respectively. Quantification estimates were imported into R/Bioconductor with tximport^48^, and differential gene expression between growth phases and temperatures was assessed with DESeq2 v1.26.0 (REF^49^) using the default values. Genes with an adjusted p-value <=0.05 were considered to demonstrate significant differential gene expression between the indicated sample groups. We report log2 fold change values for these significantly upregulated or downregulated transcripts.

### Protein extraction and proteomic analysis

After harvesting the cultures, the *T. maritima* frozen cell pellets were defrosted on ice, washed twice with 10 mL of 50 mM (Tris pH 8), and resuspended in 100 µL of the same buffer. Cell suspensions were sonicated for 20 s (x4) to lyse the cells. Cell debris and unbroken cells were removed by centrifugation at 10,000 rpm for 10 min at 4°C. The cell-free protein extracts were transferred to a 1.5 mL LoBind tube (Eppendorf) for further processing. Protein concentrations were determined with the Qubit^®^ protocol for protein quantification (ThermoFisher). Proteins (60 µg) were loaded to 12% Mini-PROTEAN® TGX™ Precast Protein Gels, 10-well, 50 µl (BIO-RAD) and run for 20 min at 120 V. Proteins were visualized by staining the gels for 3 h at room temperature with Colloidal Blue Staining Kit (ThermoFisher Scientific), washed with ultrapure (UP) water and de-stained for 18 h in UP water. Disulfide bridges were reduced with 20 mM dithiothreitol (DTT) in 50 mM ammonium bicarbonate (ABC) for 1 h at room temperature. Gels were washed with UP water (x3), followed by alkylation with 20 mM acrylamide in ABC. Each gel lane containing the samples was cut individually and sliced into smaller pieces of ca. 1 mm^2^ and transferred to 0.5 mL-Protein LoBind tubes (Eppendorf). Samples were incubated at room temperature for 15 h in 200 µl of 0.05 ng/ul of trypsin solution. The enzymatic digestion was stopped and acidified by adding 10% trifluoroacetic acid until the pH decreased between 2-4. Peptides were extracted by loading the samples onto an activated clean-up µcolumn containing two C18 Attract SPE™ (Affinisep) disks and ca. 2 mg Lichoprep RP-18 (Merck) as column material (25-40 µm). The column was washed with 100 μl of 1 ml/1 HCOOH in water and eluted with 100 µl of acetonitrile : 1ml/l formic acid in water (1:1). Samples were concentrated with an Eppendorf concentrator at 45°C for about 2 h. The volume of each sample was adjusted to 50 μl and stored at -20°C until they were injected in the nLC 1000 (Thermo EASY nLC) MSMS as described by REF^50^. The MS/MS spectra were analyzed with MaxQuant 1.5.2.8 (REFs^51,52^) with default settings for the Andromeda search engine completed by on-default variable modification settings for the deamidation on N and O. The protein sequence database for *T. maritima* MSB8 (downloaded from Uniprot March 2019) together with a contaminants custom database that contains sequences of common contaminants like Trypsins (P00760, bovin and P00761, porcin) and human keratins (Keratin K22E (P35908), Keratin K1C9 (P35527), Keratin K2C1 (P04264) and Keratin K1CI (P35527)) was used to identify the protein’s identities based on the detected peptides. The “label-free quantification” as well as the “match between runs” options were enabled. De-amidated peptides were allowed to be used for protein quantification and all other quantification settings were kept default. An intensity‐based label‐free quantification (LFQ) method^53,54^ was used for statistical comparisons (t-test) of normalized intensities of the protein groups between growth phases and temperature analysis. Filtering and statistical analyses were performed with Perseus v1.6.1. LFQ intensities were used for the analyses. For protein identification, groups were filtered to contain only proteins with at least two peptides of which at least one should be unique and at least one should be unmodified. Reverse hits and contaminants were filtered from the dataset. LFQ values were transformed to log10. For calculations, LFQ missing values were replaced by 6.8, a value slightly below the lowest measured value. A two-sample t-test with a false discovery rate (FDR) threshold set to 0.05 and significance: S0 = 0.05 were applied for comparisons. A protein was considered to be significantly up or down regulated when the protein abundance ratio was >0.05. Protein level changes between growth phases or growth temperatures were visualized with voronoi-treemaps^55^ using the Paver Software (Decodon, GmbH). The functional annotation was retrieved from the Kyoto Encyclopedia of Genes and Genomes (KEGG) for *T. maritima* MSB8. The data utilized for visualization depicts the log10-ratios of the mentioned comparison.

### Selection of potential MSL synthases

The selection criteria for the detection of MSL synthases included those genes found to be activated either in the *T. maritima* transcriptome and/or proteome, and in the *T. ethanolicus* transcriptome, respectively, following the criteria specified above. The selected genes and proteins for the MSL synthase were retrieved after screening their annotation and assigned biosynthetic pathways with KEGG and UniProt (The UniProt Consortium 2021). Searches for potential homologous proteins in *Butyrivibrio fibrisolvens* and *Clostridium ventriculi* were performed with the PSI-BLAST algorithm (Position-Specific iterated BLAST)^56^ at the protein level search using the UniProt ID from the selected radical or membrane proteins as a query (Supplementary Table 15). For *T. ethanolicus*, we performed a blast search of the candidates against NCBI NR database (downloaded November 2020) removing proteins belonging to any species from the *Thermoanaerobacter* genus (taxid: 1754). We achieved this by providing a negative sequence id list (Supplementary File 3), the search was done with a minimum e-value of 0.0001 and 50 maximum target sequences, and we kept the best hit for each query sequence.

We downloaded all genomes from species that are known to produce DAs from the PATRIC genome database^57^ on January 29th, 2021. Apart from *T. maritima*^4^, other diabolic acid producers include *Butyrivibrio fibrisolvens, Clostridium ventriculi, and Fervidobacterium pennivorans, Pseudothermotoga elfii, Pseudothermotoga hypogea, Pseudothermotoga lettingae, Thermosipho africanus, Thermosipho melanesiensis, and Thermotoga neapolitana* (Supplementary Table 21). No genomes of *Pseudothermotoga subterranea* were present in the PATRIC database. For *T. maritima*, these include the assemblies^47,58^, on which the GenBank and UniProt annotations are based, and that were used for our transcriptomic and proteomic analyses (PATRIC ID 243274.17 (GCA_000390265.1_ASM39026v1) and 243274.5, respectively).

Completeness and contamination were estimated with CheckM v1.1.3 (REF^59^) in lineage_wf mode, and genomes whose completeness – 5 x contamination was lower than 70% -- were excluded, as were plasmids. The resulting 50 genomes were annotated with Prokka v1.14.6 (REF^60^) with the --kingdom flag set to Bacteria. Orthologues were identified with Roary v13.3.0 (REF^61^), with the minimum percentage identity for blastp (-i) set to 10% and MCL inflation value (-iv) set to 1.5 to allow for sequence divergence of the highly diverse set of species, and the -s flag (do not split paralogs) set. Transmembrane helices were identified in the predicted proteins with TMHMM v2.0c (REF^62^) http://www.cbs.dtu.dk/services/TMHMM/. Proteins predicted by Prokka were linked to those in the GenBank and UniProt annotation files by bi-directional best blastp hit (the GenBank proteins to the proteins predicted from PATRIC identifier 243274.17, and the UniProt proteins to the proteins predicted from ID 243274).

### Recombinant production of candidate MSL synthases and GeR coding genes

To examine the potential enzymatic activity of the candidate MSL synthases and GeR coding genes (see Supplementary Tables 16, 17, and Supplementary Information), they were commercially synthesized (Eurofins, Germany), subcloned in pET29b or pCDFDuet-1 and expressed in the *E. coli* BL21 DE3 strain. For the case of the potential MSL synthase genes (see Supplementary Tables 16, 17) liquid cultures (50 ml) of exponentially growing *E. coli* harboring the empty expression vector or the vector including the coding gene were grown in 2×YT medium at 37°C and induced with 0.2 mM IPTG for 3 and 16 h both anaerobically and aerobically. To test the activity of the potential modified-plsA genes (GeR-coding gene), liquid cultures of exponentially growing *E. coli* harboring an empty pET29b or pET29b-coding gene were grown in 2×YT media at 37°C, induced with 0.2 mM IPTG, and incubated at 25°C for 3 h and 16 h both anaerobically and aerobically. The expression of the proteins was verified using 12% Mini-PROTEAN TGX precast gels (Bio-Rad), stained with Colloidal blue staining (Invitrogen™).

### Homology searches of MSL synthases and GeR enzymes and phylogenetics

Confirmed MSL synthase of *T. ethanolicus* and glycerol ester reductase (GeR) of *T. maritima* MSB8) were queried with DIAMOND v2.0.6 (REF^63^) against the NCBI non-redundant protein sequence database (nr) (REF^64^) downloaded on 7 January 2021. Hits with an e-value =< 1e^-50^ and query coverage of >= 30% were selected. Proteins were clustered with cd-hit v4.8.1 (REF^65^) using a sequence identity threshold of 0.7, and representative sequences were aligned with Clustal Omega v1.2.4 (REF^66^). Gaps in the alignments were removed with trimAl v1.4.rev15 [REF] in -gappyout mode. Phylogenetic trees were constructed with IQ-TREE v2.1.2 (REF^67^) with 1,000 ultrafast bootstraps. Model selection^68^ was based on nuclear models and the best-fit model was chosen according to BIC (LG+R10 for both genes). Maximum-likelihood trees were visualized in iTOL^69^.

### 3D Model and protein domain analysis

Sequences of MSL synthases of *iso*-DA and DA producers were retrieved with BLASTP^70^. The alignment was performed with MAFFT (REF^71^) in the https://www.ebi.ac.uk/Tools/msa/mafft/ server using the BLOSUM62 substitution matrix, a gap open penalty of 1.53 and a gap extension of 0.12. The multiple sequence alignment was edited with Jalview^72^ and the amino acid regions forming the conserved blocks in all MSL synthases were retrieved by coloring by 100% percentage identity of conservation between the proteins. Structure-based alignment showing conserved hydrophobic regions^73^ between *iso*-DA and DA producers were colored based on percentage of conservation (90%) of the hydrophobic or hydrophilic residues. The secondary structure prediction showed in the alignment was performed with Jpred. The 3D models of the proteins were calculated with AlphaFold V2.1.0 with the Google Colab platform (https://colab.research.google.com/github/deepmind/alphafold/blob/main/notebooks/AlphaFold.ipynb), accessed in February 2022, with no templates and refined using the relax option. The resulting prediction was visualized using the PyMol software^74^.

## Data availability

RNAseq and proteomics data is in the process of being submitted to NCBI and ProteomeCentral, respectively. Lipid analysis raw files will be deposited in Zenodo.org upon acceptance of the manuscript. All raw materials are available to reviewers upon request.

## Code availability

Not applicable

## Acknowledgements

We thank Marcel van der Meer for help in the labeled incubation data interpretation. This project received funding from the European Research Council (ERC) under the European Union’s Horizon 2020 research and innovation program (grant agreement no. 694569-MICROLIPIDS) to JSSD. LV and JSSD receive funding from the Soehngen Institute for Anaerobic Microbiology (SIAM) through a Gravitation Grant (024.002.002) from the Dutch Ministry of Education, Culture, and Science (OCW). KF (PI-LV) receives funding from the Simons-Moore foundation.

## Contributions

DXSC designed the experiments, executed the experiments, analyzed the data, interpreted the results and wrote the paper; MS designed and executed part of the experiments, JCE, AAA contributed to the bioinformatic analysis, SB executed and interpreted the proteomic data, MK and NB performed the lipid analysis and interpretation, FABvM contributed to the bioinformatic and phylogenetic analyses, LSvS and KF contributed to the gene expression assays. JSSD and LV acquired funding, contributed to the design of the experiments, interpretation of the data, and writing of the manuscript. All co-authors have read and approved the manuscript.

