## Supplemental Information for "Disentangling the lipid divide: Identification of key enzymes for the biosynthesis of unusual Membrane-spanning and Ether lipids in Bacteria"

### Supplementary Information

The Supplementary Information includes supplementary results and discussion of the main text and it is organized in the same subsections. In this context, it is relevant to note that Supplementary Figure 1 includes the chemical structures of diabolic acids (DAs), *iso*-diabolic acid (*iso*-DA), glycerol ether derivative of *iso*-diabolic acid, and branched glyceryl dialkyl glycerol tetraethers (brGDGTs) as discussed in the text. Figure 1a, includes the coupling reactions of iso-fatty acids (FA) to form iso-diabolic acid via tail-to-tail condensation at the ω positions, and of saturated FA to form diabolic acid via tail-to-tail condensation at the ω-1 positions.

#### *Iso*-diabolic acid is produced via condensation of *iso*-fatty acids

##### *1) Synthesis of MSLs in the iso-diabolic acid producer* **Thermoanaerobacter ethanolicus**

We investigated the composition of the core membrane lipid (i.e., after removal of the polar head group by hydrolysis) of *T. ethanolicus* across growth phases (early exponential, mid-exponential, late-exponential, and stationary phases) at both 60°C (optimal growth temperature) and 45°C (suboptimal temperature).

At 60°C, the core lipids detected were mainly composed of $C_{15:0}$ and $C_{17:0}$ *iso* and *anteiso* FAs, *n*-$C_{16:0}$ FA, and the $C_{30:0}$ *iso*-DA (see Supplementary Table 1). A range of other FAs and $C_{31}$ and $C_{32}$ *iso*-DAs and 13-methyl $C_{28}$ (total of 29 carbon atoms) and 13-methyl $C_{30}$ (total of 31 carbon atoms) diacids were detected in smaller relative abundances. Various, predominantly methyl-branched, hydroxy (OH) FAs ($C_{15:0}$ to $C_{18:0}$) were also detected. Dimethyl acetals (DMA), which are derived from vinyl ethers upon acid hydrolysis, were also present in small amounts with *iso*-$C_{15:0}$, *iso*-$C_{16:0}$, *n*-$C_{16:0}$, *iso*-$C_{17:0}$ and *anteiso*-$C_{17:0}$ carbon skeletons.

The relative distribution of specific FAs and *iso*-DAs changed significantly across growth phases as observed for the dominant *iso*-$C_{15:0}$ FA, which comprised 51% during the early exponential phase, increased to 55% in the mid-exponential phase and decreased to 46% at the late-exponential phase, finally representing 38% of the core lipids detected at the stationary phase of growth (Figure 1b, Supplementary Table 1). Similarly, other minor FAs decreased in their abundance across growth phases, the $C_{16:0}$ (from 7.8% to 3.4%) and the *iso*-$C_{17:0}$ (from 10.0 to 4.6%). In contrast, a marked increase in the $C_{30}$ *iso*-DA was detected during the growth experiment; it represented 4.5% at the early exponential, slightly decreased in the mid-exponential phase to 4.3%, increased to 15.4% during the late-exponential phase, and finally represented a 24% at the stationary phase. Some other FAs and *iso*-DAs also showed a minor increase during the time-course growth experiment; the *iso*-$C_{13:0}$ increased 2.5% in relative abundance, the *anteiso*-$C_{15:0}$ 1.7%, the *iso*-1-OH-$C_{15:0}$ 2.6%, and the 13-methyl-$C_{28}$ diacid 1.0% from the early-exponential to the stationary phase of growth. Overall, a significant decrease (ca. 18.2%) of the *iso*-$C_{15:0,}$ and *iso*-$C_{17:0}$ coincided with the increase (approx. 20%) on the major $C_{30}$ *iso*-DA and 13-methyl-$C_{28}$ diacid.

At a growth temperature of 45°C, a similar range of FAs, methyl-branched FAs, diacids and *iso*-DAs, OH FAs, and DMAs, as observed at 60°C were detected (Supplementary Table 1). The relative abundance of specific FAs and *iso*-DAs changed significantly across growth phases as observed for the dominant *iso*-$C_{15:0}$ FA, which relative abundance amounted 48% during the early exponential phase, increased to 61-62% in the mid- and late-exponential phases, and finally represented 50% at the stationary phase of growth. Similarly, other minor FAs decreased in their abundance across growth phases, the $C_{16:0}$ (from 7.1% to 1.0%), the *anteiso*-$C_{15:0}$ (from 11 to 3.8%) and the *iso*-$C_{17:0}$ (from 7.2 to 3.6%). In contrast, an increase in the $C_{30}$ *iso*-DA was evident; it represented a 1% at the early and mid-exponential phase, slightly increased to 2% during the late-exponential phase, and represented a 14% at the stationary phase. Some other FAs also showed a minor increase during the time-course growth experiment; the *iso*-$C_{13:0}$ increased 4.3% in relative abundance,  and the *iso*-1-OH-$C_{15:0}$ increased 6% from the early-exponential to the stationary phase of growth. Overall, a significant decrease (ca. 8.9%) on the *iso*-$C_{15:0,}$ *anteiso*-$C_{15:0}$ and *iso*-$C_{17:0}$ coincided with the increase (ca. 12.8%) on the $C_{30}$ *iso*-DA (Supplementary Table 1).

In addition, we performed experiments adding labeled branched amino acid ($^{13}$C-leucine), precursor of *iso*-branched chain FAs, to *T. ethanolicus* cultures. Results are summarized in the main text in Figure 1b and in Supplementary Table 2.

##### *2) Synthesis of membrane-spanning lipids in the diabolic acid producer* **Thermotoga maritima**

The lipid composition of the cultures of *T. maritima* across different growth phases and temperatures is included in Figure 1b, Supplementary Figure 2, and Supplementary Table 3.

#### In search for potential proteins for the biosynthesis of membrane-spanning and ether lipids in bacteria

Our results with *T. ethanolicus* indicate that the biosynthetic reaction leading to *iso*-DA is growth phase-dependent (Figure 1c, Supplementary Figure 2), as also shown for the formation of the MSL DA in *T. maritima* (Figure 1b, Supplementary Table 3). In both *T. ethanolicus* and *T. maritima*, the percentage of MSLs substantially increased during the stationary phase of growth, suggesting that experimental conditions allow for the detection of the activation of the genes coding for the proteins involved in the synthesis of MSLs. To test this, we analyzed the transcriptomic and proteomic response of these two bacterial species and compared them between different growth phases and at optimal and suboptimal growth temperatures.

**1) *Changes in gene regulation of* Thermoanaerobacter ethanolicus *across growth phases***

The genome of *T. ethanolicus* is composed of 2,876 genes and 67 RNA genes, 2638 are protein-coding genes and 171 pseudogenes[1]. At the optimal temperature of growth (60°C) and across growth phases, 69–81% of the 2,876 genes were found significantly regulated at the transcriptomic level; 35–42% of these genes were found to be upregulated (i.e., an 1.5 fold change). The experiment at suboptimal growth temperature (45°C) and across growth phases showed between 41−77% of the 2,876 genes to be significantly regulated at the transcriptomic level; 21−39% of these genes were upregulated. These numbers are comparable to those found at optimal growth temperature (i.e., 60°C).

To be able to distinguish between the most significant changes across the transcriptome of *T. ethanolicus*, a hierarchical representation from the annotated genome mainly based on the KEGG pathways database[2], and using Voronoi trees[3] was applied for visualization of the changes and the results as discussed below.

During the mid to exponential phase at the optimal growth temperature (60°C), upregulated genes (Supplementary Table 4) were those involved in the metabolism of amino acids (histidine metabolism), two-component system, the ABC transporters, and the peptidoglycan biosynthesis (Supplementary Figure 3). Similarly, during this growth phase at suboptimal temperature conditions (45°C) the main upregulated metabolic pathways detected were those involved in the biosynthesis of amino acids, metabolism of terpenoids and polyketides, and processes related to cell motility (Supplementary Figure 3). At both temperatures, the histidine metabolism was found significantly upregulated, suggesting that during the early growth phases histidine might be specifically required for the proper growth and adaptation.

During the late-exponential phase to the stationary phase at optimal growth temperature, the genes involved in DNA replication and repair, biosynthesis of secondary metabolites were found significantly upregulated (Supplementary Figure 3). At suboptimal temperature conditions, the main enriched categories were membrane transport and D-amino acid metabolism (Supplementary Figure 3). At both conditions of growth, the galactose metabolism and the uptake system the phosphoenolpyruvate carbohydrate phosphotransferase system (PTS) was found to be upregulated, indicating that the sugar transport system is active and required for growth and the synthesis of other compounds in the cell.

During the mid-exponential to stationary phase, major changes resulted in the activation of membrane transport, such as the ABC transporters, nucleotide metabolism, and biosynthesis of amino acids. The metabolism of galactose and fructose was also activated. At suboptimal growth temperature, membrane transport, galactose metabolism, the PTS system and the histidine metabolism were upregulated.

**2) *Changes in gene regulation of* Thermoanaerobacter ethanolicus *between different growth temperatures***

Comparison between growth temperatures were examined specifically between growth phases. Supplementary Table 5 shows the average expression at each growth phase of all the upregulated genes at optimal growth temperature when compared to suboptimal growth temperatures. From the transcriptomic analysis, we detected 1,410 significantly regulated genes at the mid exponential growth phase, 1,949 during the late exponential phase and 2,106 at the stationary phase (Supplementary Table 5). Similarly, as in the growth phase analysis, genes were clustered by their predicted metabolic pathways by KEGG classification and visualized with Voronoi-tree maps. This analysis revealed that the metabolic classification at the mid-exponential growth phase that was upregulated was related to the KEGG categories genetic information processing, metabolism of other amino acids, and cell motility (Supplementary Figure 4). During the late-exponential phase, membrane transport, metabolism of other amino acids and cellular processes were upregulated (Supplementary Figure 4). Finally, during the stationary phase, the metabolic pathways that were upregulated were those involved in environmental adaptation, cellular processes, replication and repair, and the biosynthesis of other secondary metabolites (Supplementary Figure 4).

Our analysis indicated that among the KEGG categories that grouped enzymes whose genes were more upregulated during all growth phases, the most relevant was that of metabolism of other amino acids including, D-amino acid metabolism. The synthesis of D-amino acids has been associated with different processes of adaptation in Bacteria such as biofilm development[4], growth fitness[5], etc. D-amino acids are a structural part of the bacterial cell wall as they are incorporated into peptidoglycan and regulate the composition, strength, and structure of the bacterial cell wall[6]. Thus, the synthesis of D-amino acids may be required as an adaptation for growing at high temperatures for this organism or as previously proposed[7], a common strategy of bacteria for adapting to non-optimal environments.

When compared to the suboptimal temperature of growth, lysine degradation was upregulated at increased temperatures during all growth phases, indicating an active amino acid utilization. Lysine has been demonstrated to be related to adaptation to environmental stresses (i.e., osmotic stress, oxidative stress). It is possible that the products of lysine degradation are involved in the adaptability of the cell to high-temperature conditions. In this analysis, the citrate cycle (TCA) was also upregulated during all growth phases at a high temperature of growth, thus indicating greater energy demands when cells are grown under these conditions. To our knowledge, a temperature-dependent analysis of what? has never been reported for *T. ethanolicus*. Proteomic profiling of a related species *Thermoanaerobacter tengcongensis*[8] identified increased temperature-dependent proteins related to glycolysis. In contrast, we only detected the upregulation of glyceraldehyde-3-phosphate dehydrogenase (WP_003870200.1_2183) during the mid- and late-exponential phases, while the other glycolysis-related proteins; oxaloacetate decarboxylase (WP_003870653.1_1774) and dihydrolipoyl dehydrogenase (WP_003870815.1_712) were found to be downregulated, and only the dihydrolipoyl dehydrogenase was upregulated during the stationary phase. This shows that energy production through glycolysis proteins is not activated during all growth phases, nor in all *Thermoanaerobacter* related species.

In addition, during growth at elevated temperatures, the chaperonins, GroES and GroEL, were found to increase in *T. tengcongensis*[7]. Similar results were found in *T. ethanolicus*, the GroEL chaperonin (WP_003870513.1_1078) was also upregulated during late-exponential and in minor abundance during the stationary phase. Additionally, the molecular chaperones Hsp33 (WP_003870176.1) and Hsp20 (WP_003870450.1_230) were upregulated during the late-exponential phase. While the Hsp33 and Hsp70 (WP_003871477.1_2461) were upregulated during the stationary phase

**3) *Activation of lipid metabolic pathways during growth phases at optimal and suboptimal growth conditions in* Thermoanaerobacter ethanolicus**

At optimal temperature conditions, the gene coding for the glycerol kinase (WP_003871378.1) *glpK*, which converts glycerol to phosphatidic acid (PA), was upregulated at all examined growth phases, (Supplementary Table 6, Supplementary Figure 5). The gene *plsX*, coding for acyl-ACP acyltransferase (WP_003869200.1), which catalyzes the conversion of acyl-ACP and inorganic phosphate (Pi) to acyl-phosphate, was upregulated during the late exponential phase to stationary transition. The *plsY* gene, coding for the enzyme that transfers acyl-phosphate (acyl-PO4) to G3P leading to 1-acyl-G3P; the acyl phosphate-G3P-acyltransferase protein (WP_003868555.1), was also upregulated during the mid-exponential to the late exponential phase at both, optimal and suboptimal growth conditions (Supplementary Table 6, Supplementary Figure 5). During all the examined growth phases, and at both growth temperatures, the enzyme involved in the second acylation step of the glycerol backbone, the 1-acyl-*sn*-G3P acyltransferase (WP_003870115.1/WP_003870040.1), encoded by the *plsC* gene, was upregulated. The enzyme that activates PA for the phospholipid head synthesis, the cytidine diphosphate (CDP)-diglyceride synthase (WP_003869142.1), encoded by *cdsA*, was upregulated only at suboptimal growth temperature, during the mid-exponential to late-exponential phases. Finally, the *pgsA* gene coding for a CDP-diacylglycerol (DAG) G3P 3-phosphatidyltransferase (WP_003869119.1/ WP_003870898.1), which catalyzes the conversion of CDP-DAG to phosphatidyl-glycerolphosphate (PGP) or the anticipated intermediate for the phosphatidylglycerol (PG) phospholipid, was upregulated at both, optimal and suboptimal temperature of growth during all the analyzed growth stages (Supplementary Table 6, Supplementary Figure 5). In general, the transcriptomic analysis of *T. ethanolicus* at different growth phases supports the activation of the lipid biosynthetic pathways leading to the formation of phospholipids as intermediates to the formation of iso-DA (Supplementary Figure 5).

**4)** ***Changes in gene regulation of*** **Thermotoga maritima** ***across growth phases***

The genome of *T. maritima* is composed of 1,904 genes and 52 RNA genes, 1,852 are protein-coding genes and 15 pseudogenes[1]. At optimal temperature of growth and across growth phases, between 25 to 67% of the 1,904 genes (including RNA genes) were found to be significantly regulated at the transcriptomic level. From these, 14−34% of these genes were upregulated. At the proteomic level, 21−38% of the 1,852 proteins were found significantly regulated, from which 10−22% were upregulated.

The experiment at suboptimal temperature (55°C) and across growth phases showed that 41 - 65% of the 1,904 genes (including RNA genes) were significantly regulated at the transcriptomic level. From the significantly regulated genes, 18–33% of the genes were upregulated, comparable to the results obtained at optimal growth conditions. At the suboptimal temperature, the transcriptomic analysis between the early and stationary growth phase and between exponential and stationary growth phase showed that RNA genes were mostly downregulated. At the proteomic level, between 12−31% of the 1,852 protein-coding genes were found significantly regulated, from which 7 - 15% were upregulated. Between the exponential and stationary phase of growth, the changes in protein abundance were not statistically significant with respect to the threshold (P-value 0.05).

**4a) From the early exponential phase to the exponential phase**. In the early to exponential phase at the optimal growth temperature, 523 genes were upregulated (Supplementary Table 7) from which the main KEGG categories represented at a transcriptomic level were those related to energy metabolism, cellular processes such as cell motility, and genes involved in signal transduction (Supplementary Figure 6a). Similar findings were detected during this growth phase at suboptimal temperature conditions, from 361 upregulated genes (Supplementary Table 8); the main enriched categories were those related to cell motility and signal transduction (Supplementary Figure 6b). At the transcriptomic level, the processes that were significantly upregulated at both growth temperatures were those related to cell motility, pointing to an increase in motility during the early growth phases. The upregulation of these genes would allow the bacterial cell to increase its motility during unfavorable environmental conditions or at the exponential phase.

The proteomics analysis at optimal growth temperature detected 312 upregulated proteins (Supplementary Table 9), mainly belonging to those involved in the biosynthesis of secondary metabolites, terpenoids and polyketides (bioactive natural products), co-factors and vitamins, and other amino acids (Supplementary Figure 7a). On the other hand, proteomic analysis of the *T. maritima* cells grown at suboptimal temperatures detected 136 upregulated proteins (Supplementary Table 10), which were mostly related to processes involved in cell motility, signaling, and cellular processes and proteins implicated with the metabolism of terpenoids and polyketides, together with energy metabolism (Supplementary Figure 7b). The detected upregulation of proteins involved in the biosynthesis of terpenoids and polyketides at both conditions is compatible with the production of specific metabolites, which might be essential for survival during early-growth phases; although often classified as secondary metabolites, their production is often associated with providing an advantage to the bacteria to a specific ecological niche. On the other hand, at optimal temperature conditions, the observed upregulation of proteins involved in the synthesis of co-factors and vitamins points to as being essential for growth. Furthermore, the proteomic analysis at suboptimal temperature conditions also showed increased upregulation of proteins related to cell motility, as also observed at the transcriptomic level, which further supports that cells growing at a lower temperature than the optimal have the tendency to be more motile.

**4b) Changes from the exponential phase to the stationary phase.** During the transition from the exponential to the stationary phase, 275 genes were upregulated (Supplementary Table 7), which are mostly related to anti-microbial drug resistance and genetic information processing like replication and repair, translation, and glycan biosynthesis (Supplementary Figure 6a). At sub-optimal temperature growth conditions, 511 upregulated genes (Supplementary Table 8) involved in cellular processes like cell motility, genetic information processing such as transcription, replication and repair, signal transduction, and cellular communities of prokaryotes, were detected (Supplementary Figure 6b). Genes involved in anti-microbial drug resistance are mainly related to biosynthetic pathways producing genes encoding proteins for the synthesis of various antibiotic agents (i.e., cationic antimicrobial peptides (CAMP), β-lactam and vancomycin resistance in

*Thermotoga*[2]. *T. maritima* is known to be naturally resistant to aminoglycosides[9,10]; therefore, the upregulation of these genes could imply the activation and synthesis of the proteins involved in CAMPs. Genes involved in β-lactams and vancomycin drug resistance are also activated through the stationary phase at optimal growth conditions potentially providing resistance to these antibiotics.

In addition, at both growth temperatures, the activation of genetic processes such as replication and repair suggest that mechanisms involved in mismatch repair, nucleotide excision and repair, homologous recombination and DNA replication, or the activation of mechanisms to recognize and repair DNA damage are important for preparing cells for the stationary phase and growing at optimal (high) temperature. This is in line with the high levels of homologous recombination previously observed in *Thermotoga* sp.[11], a process that is thought to be needed for DNA repair in thermophiles[12,13]. On the other hand, the biosynthesis of the cell envelope was found differentially activated at optimal temperature of growth, specifically the glycan biosynthetic pathway, indicating that during optimal growth conditions, the cell is actively forming the cell wall for the stationary phase. As detected during early to the exponential growth phase, processes related to cell motility were also upregulated during growth at suboptimal temperature, suggesting an increase in motility on the cells grown at this condition and across all the examined growth phases. As stated earlier, the upregulation of motility genes would allow the bacterial cell to increase its motility during unfavorable suboptimal (colder) temperature conditions.

At the proteomic level, 275 proteins (Supplementary Table 9) were upregulated at the optimal growth temperature, mainly those involved in the biosynthesis of other secondary metabolites as antibiotics, the metabolism of terpenoids and polyketides, membrane transport, and the antimicrobial drug resistance (Supplementary Figure 6a). At sub-optimal temperature conditions, no proteins were statistically significantly upregulated. Nevertheless, with the Voronoi treemap analysis we identified some minor changes mainly in the metabolism of amino acids, carbohydrates, co-factors, and vitamins, but also some in some hypothetical proteins (Supplementary Figure 6b). The activation of the biosynthesis of other secondary metabolites during growth at optimal temperature, which includes novobiocin and monobactam antibiotics together with the biosynthesis of terpenoids and polyketides, indicates that the production of secondary metabolites may be necessary for the cells in the stationary phase, as these compounds possess specific biological activity. In addition, membrane transport activation during the transition to the stationary phase could be attributed to the capability of *T. maritima* to efficiently metabolize a variety of carbon sources or other metabolites that might serve to acquire nutrients at the stationary phase and for survival.

**4c) General changes from the early exponential phase to the stationary phase.** At the transcriptomic level at this growth phase transition, 635 upregulated genes (Supplementary Table 7) were detected, which are mainly involved in environmental information processing such as signal transduction, cellular processes, and cell motility classification. Also, genes involved in the metabolism of terpenoids, polyketides and energy metabolism, genetic information processing as replication and repair were significantly upregulated (Supplementary Figure 6a). At suboptimal temperature, 652 upregulated transcripts (Supplementary Table 8) were detected. They are involved in cellular processes of cell motility, signal transduction, cellular community prokaryotes, metabolism of terpenoids and polyketides, genetic information processing as replication and repair and also some belong to unclassified metabolism classification (Supplementary Figure 6b).

The analysis of the major changes at the transcriptomic level revealed that at optimal growth temperature, the main upregulated classification was signal transduction, which enables the bacterial cell to sense, respond, and to adapt to changes in their environment or in their intracellular state, or detect changes in cell physiology through the two-component systems, later activating and regulating gene expression. The activation of these signaling systems likely indicates that cells can adapt to changing conditions by efficiently producing the needed metabolites to survive the growth conditions. Besides, as seen at both evaluated temperatures, an upregulation of processes of the cellular community of prokaryotes classification (quorum sensing) was detected. Quorum sensing is known to be driven by changes in cell-population density, and, therefore, it suggests that the cells are able to detect variation in cell number during growth and subsequently coordinate social functions such as biofilm formation, competence, among others. It is possible that as cell density increased during the growth phase transition, this could be detected through this system in *Thermotoga*, thus allowing cells to regulate their physiological responses. Also, as observed between the transition of the early exponential and the exponential phase, cellular processes like cell motility were also found upregulated, indicating an increase in motility not only during the early growth phases but also during the overall transition from early growth to the stationary phase.

The metabolism of terpenoids and polyketides, specifically the terpenoid backbone biosynthesis, producing the ansamycin antibiotic and carotenoids, suggest these compounds may contribute to growth and stability to the cell along the growth cycle.

At the proteomic level, our analysis indicated that 293 of the upregulated proteins (Supplementary Table 9) were classified as metabolic proteins, including those within the categories of xenobiotics metabolism and biodegradation, metabolism of terpenoids and polyketides, other amino acids, cofactors and vitamins, nucleotide, and biosynthesis of other secondary metabolites, as well as unclassified metabolism. Also, proteins that are not included in the pathway or that are poorly characterized were enriched in this analysis (Supplementary Figure 6a). During growth at suboptimal temperature, at proteomic level, 411 proteins were upregulated (Supplementary Table 10), belonging to the xenobiotic biodegradation and metabolism, and the metabolism of cofactors and vitamins, amino acids, and energy classification (Supplementary Figure 6b). At both growth temperatures, activation of xenobiotic degradation was detected, which includes nitrotoluene, chloroalkane, naphthalene and benzoate; compounds that are not usually present on the growth media, neither reported to be produced by this organism, it is possible that the enzymes involved in the degradation of this compounds are using different substrates from the media, or alternatively that similar compounds could be formed as by-products from growth and became toxic for *Thermotoga* cells, thus activating this common mechanism for degradation at both temperature conditions. Similarly, at both examined temperatures, the metabolism of cofactors and vitamins indicates their production might be essential for supporting growth.

**5) *Changes in gene regulation of* Thermotoga maritima *between different growth temperatures***

The differences between growth temperatures were examined specifically. The genes and proteins that were differentially upregulated (80°C *vs* 55°C) were attributed to those changes that permitted adaptation or growth at the optimal temperature (80°C). Supplementary Tables 11 and 12 show the average expression of all the upregulated genes and proteins that are reported to be involved in thermoregulation at optimal growth temperature when compared to suboptimal growth temperatures for each growth phase. From the transcriptomic analysis, we detected 391 upregulated genes in the early growth phase, 450 during the exponential phase and 593 in the stationary phase (Supplementary Table 11).

At the transcriptomic level, at the early growth phase, the upregulated transcripts were mostly classified in the KEGG categories of glycan degradation, metabolism of other amino acids, starch and sucrose metabolism, and quorum sensing (Supplementary Figure 7). At the proteomic level, the riboflavin metabolism, the biosynthesis of amino acids, the nicotinate and nicotinamide metabolism and the pantothenate and CoA biosynthesis were activated. The activation of nicotinate and nicotinamide metabolism could indicate increased demand for co-enzyme precursors involved in redox reactions and indirectly into other biosynthetic pathways for growth at high temperature.

In both the transcriptomic and proteomic analyses, the C5-branched dibasic acid metabolism was activated, suggesting that several organic acids could be produced trough this pathway and were required for adaptation at high temperatures. At the exponential growth phase, the metabolic pathways that were upregulated in the transcriptomic analysis were the metabolisms of other amino acids, starch and sucrose metabolism, ABC transporters, membrane trafficking, and the nicotinate and nicotinamide metabolism. The activation of membrane trafficking or ABC transporters during the exponential phase indicates that microbial cells are actively translocating the substrates across membranes and, therefore, might be implicated in an extensive range of processes.

The proteomic analysis at the exponential growth phase, showed the upregulation on the biosynthesis of amino acids, prenyltransferases, and the pathway involved in the transfer of RNA and biogenesis (Supplementary Figure 7). The activation of prenyltransferases suggest there is a demand on the production of polyprenyl compounds, likely involved in adaptation to high temperature. Finally, during the stationary phase, the metabolic pathways that were upregulated in the transcriptomic analysis were the biosynthesis of amino acids, the metabolism of other amino acids, starch and sucrose, the C5-branched dibasic acid metabolism, and the membrane transport. At the proteomic level the biosynthesis of amino acids, transfer RNA and biogenesis, also the prenyltransferases and terpenoid metabolism were found upregulated (Supplementary Figure 7).

Temperature-dependent analysis at genomic and proteomic level had been previously reported on *T. maritima*[13], highlighting the activation of enzymes participating in central carbohydrate metabolism or other features related to thriving at high temperatures such as thermostable enzymes and proteins in *Thermotoga*[14]. Genomic analysis also led to the prediction of genes coding for heat shock proteins (dnaJ/K, groEL/ES, grpE, hslU/V and hsp) or cold shock proteins (cspB/L) encoded in the genome of *T. maritima*[15]. However, these studies have either covered the genomic, transcriptomic or proteomic aspect or have only focused on the specific genes or protein targets already known to contribute to the adaptation to temperature. In the present work, we extended and combined multiple analytical approaches, which led us to identify which are the major metabolic and genetic processes occurring during growth at high (optimal) growth temperatures and across the growth phases. In addition, we also compared the expression profile with growth at suboptimal temperatures (55°C). From this, we found that during early and exponential growth phases, when compared to the sub-optimal temperature of growth, it has consistently shown upregulation of amino acid biosynthesis, carbohydrate metabolism (C5 branched dibasic acid) as well as starch and sucrose metabolism. The protein primary structure is formed by a polypeptide chain of amino acids, which intrinsically determine the structure of the proteins. Comparison of amino acids composition between thermophilic proteins has shown that charged amino acids (D, E, K, R) might be enriched (i.e. CvP bias, difference between proportions of charged and polar non-charged amino acids)[16], and the overrepresentation of I, V, Y, W, R, E and L amino acids correlated with optimal growth temperature[17]. Calculation of the CvP bias and the amino acid compositional features in proteins from *Thermotoga* species has shown a unimodal distribution, confirming the correlation with optimal growth temperatures[18]. In this sense, the biosynthetic pathways for arginine and lysine from the CvP bias as well as isoleucine, valine, tyrosine, tryptophan, arginine and leucine were found upregulated in our transcriptomic and proteomic analysis (Supplementary Tables 11, 12).

We also detected upregulation of specific genes and proteins known to be involved in thermoregulation (Supplementary Table 13); a 33kDa chaperonin (Tmari 1401), the chaperon protein DnaK (Tmari 1814), the Heme Chaperone HemW (Tmari 1173), a Lon protease (Tmari 1642), a ribosome-associated heat shock protein (Tmari 0221), and the trigger factor folding chaperone (Tmari 0694). Surprisingly, the cold shock proteins CspA and CspG (Tmari 1889 and Tmari 1691) were also found upregulated across growth phases. The adaptation of cells to high temperatures due to chaperones aid in proper folding, allowing protein stability is well known[19]; in our analysis, upregulation of the 33 kDa Chaperonin, DnaK, the heme chaperone HemW, and the trigger factor folding chaperone, are expected to be constitutive expressed in thermophiles as part of their temperature adaptation. Additionally, we have detected the activation of the CspA and

CspG cold shock proteins, suggesting their role is not limited to cold temperature conditions[20], but permits growth and adaptation to high temperature as an additional function.

**6)** ***Activation of lipid metabolic pathways during growth phases at optimal and suboptimal growth conditions in*** **Thermotoga maritima**

During growth of *T. maritima* at optimal temperature conditions, between the early exponential and the exponential growth phase, the gene coding for the enzyme producing the key intermediate forming the backbone of all glycerophospholipid molecules (G3P) the G3P-dehydrogenase GpsA (Tmari 0376) was found to be upregulated at transcriptomic level (Supplementary Table 14, Supplementary Figure 8). Similarly, at the proteomic level also the GpsA protein was upregulated, together with two key enzymes from the type II FA synthesis system of bacteria, the enoyl-acyl carrier protein (ACP) reductase FabK (Tmari 0801), and the 3-oxoacyl ACP reductase FabG (Tmari 1176) (Supplementary Table 14, Supplementary Figure 8). At suboptimal temperature in the same analyzed growth interval, we found the *gpsA* gene activated, while no other significant changes at protein level were found (Supplementary Table 12, Supplementary Figure 8).

During the transition from the exponential growth phase to the stationary phase at optimal temperature conditions and from the early exponential to the stationary at suboptimal temperature, the acyl-ACP acyltransferase PlsX (Tmari 0147) was upregulated (Supplementary Table 14, Supplementary Figure 8). Also, the glycerol-1-phosphate dehydrogenase AraM (Tmari 0283), which might be involved in forming a G1P intermediate that is not the glycerol backbone present in *Thermotoga* lipids, was also found to be activated at the transcriptomic level. At the proteomic level and between the exponential to the stationary phases, the 1-acyl-sn-G3P PlsC (Tmari 1701), was upregulated. At suboptimal conditions, we detected changes at the transcriptomic level in the *plsY* gene coding for acyl phosphate-G3P-acyltransferase (Tmari 1289), in the CDP-diglyceride synthase *cdsA* gene (Tmari 1404), as well as in the *pgsA* gene (Tmari 1876) producing the PGP intermediate for the PG phospholipid (Supplementary Table 14, Supplementary Figure 8). At proteomic level, no significant changes were found in the lipid metabolic pathway at suboptimal temperatures between the exponential or early exponential and stationary phase of growth (Supplementary Table 14). Finally, between the early exponential and the stationary phase of growth and at optimal conditions, a gene coding for a glycerol kinase *glpk* (Tmari 0954) converting glycerol to phosphatidic acid (PA) was activated; in the same growth phase but at suboptimal growth temperature the *pgsA* gene was upregulated (Supplementary Table 14, Supplementary Figure 8). In the present analysis, we detected the activation of the step leading to the glycerolipid backbone G3P being upregulated during growth phases (Tmari 0376, GpsA, Supplementary Table 14), at both optimal and suboptimal temperature of growth and with both experimental approaches (transcriptome and proteome).

Previous lipidomic analyses of *T. maritima* showed a variety of head groups in the intact polar lipids (i.e., core lipids with attached head groups; IPLs) forming the membrane, but only the phosphatidylglycerol (PG) headgroup-bearing IPLs contained MSLs[21]. Accordingly, a biosynthetic pathway for MSLs through the synthesis of PG headgroup-containing intermediates was proposed. This hypothesis is now confirmed by the upregulation of the genes coding the biosynthetic steps leading to PG lipids (Supplementary Table 14).

##### 7) *Selection of the potential membrane-spanning lipid synthases*

MSL synthase are expected to mediate the dimerization of FA building blocks to form the *iso*-DA or DA building blocks. This reaction would involve a radical intermediate formed by a hydrogen extraction at the tail of one of the fatty acids, followed by a condensation with the other tail, involving the loss of another hydrogen, resulting in the formation of a C-C bond in the absence of an activated intermediate. Such unique reactions are commonly catalyzed by radical proteins, a group that shares an unusual Fe-S cluster associated with the generation of a free radical by the reduction of S-adenosylmethionine (SAM). A conserved cysteine-rich motif (CxxxCxxC) is present in all identified radical-SAM proteins[22], which catalyze a wide variety of reactions, including unusual methylations, isomerization, and ring formation. Based on these considerations, we defined selection criteria for the detection of potential MSL synthases in the pool of genes found to be activated either in the *T. maritima* transcriptome or proteome, or in the *T. ethanolicus* transcriptome, but not yet attributed to any known metabolic pathway. These criteria were that the gene or protein (1) should code for a radical-SAM protein that contained the cysteine-rich motif, (2) should encode for an oxidoreductase utilizing an [Fe4 4S] cluster, which can act on CH or $CH_2$ groups, (3) should encode a membrane-bound protein, as most of the proteins known to be involved in the formation of core membrane lipids are membrane-associated, and (4) should be homologous to proteins in other MSL producing bacteria (Supplementary Figure 9-10, Supplementary Tables 15-19).

Our main focus for selecting the potential *iso*-DA synthases was based on the upregulation of those genes that code for proteins that were predicted to possess a radical-SAM signature, and that they would be upregulated across growth phases and at optimal temperature (Supplementary Figure 9). We performed a BLAST (pBLAST) analysis of the pre-selected (radical) SAM-coding genes upregulated in *T. ethanolicus* (Supplementary Table 15) against the NCBI non-redundant protein database excluding the *Thermoanaerobacter* genus hits in order to evaluate the presence of genes encoding similar proteins in other organisms potentially producing membrane-spanning lipids. As a result, a total of ten proteins were chosen based on their predicted radical SAM domain annotation and their distribution among other potential MSL producers; EGD52722, EGD51308, EGD51168, EGD50870, EGD50873, EGD50888, EGD50779, EGD50325, WP_129545148.1 and EGD51024 (Supplementary Table 16). The EGD50870 gene, annotated as a B12-binding domain-containing radical SAM protein and EGD50779, annotated as a radical SAM protein, possess homologous genes in organisms belonging to the order of Thermotogales producing the MSL DA. Similarly, the EGD50873, EGD51024 and EGD50888, which are annotated as radical proteins, possess similar proteins in Archaea and Clostridia. Finally, for the EGD52722 annotated as a Rossmann-like protein, EDG51308, annotated as FAD-dependent oxidoreductase, EGD51168, EDG50325 and WP_129545148.1, annotated as radical SAM proteins, we found similar proteins mainly in Clostridia. In the case of WP_129545148.1, which contains 446 amino acids (aa), and belongs to the obsolete version of the annotated genome, we noticed that the coding sequence is annotated as two different proteins (EGD51859.1 of 220 aa and EGD51860.1 of 226 aa) in the available version of the genome[23]. Based on motif analysis (Pfam) we decided that WP_129545148.1 was the correct protein (Supplementary Figure 10).

For the case of *T. maritima*, genes coding for radical SAM proteins in the transcriptome or for radical proteins annotated in the proteome of *T. maritima* and not yet attributed to any other metabolic pathway were further selected (Supplementary Figure 9). Within those, the genes coding for radical-activating enzymes that contained the cysteine-rich motif were selected: Tmari 0106, Tmari 0180, Tmari 0771, Tmari 0827, Tmari 0926, Tmari 1324, Tmari 1545 and Tmari 1609 (Supplementary Table 17). Similarly, enzymes performing this reaction could be annotated as oxidoreductases, utilizing the Fe4 4S cluster, and depending on their classification, can act on CH or CH2 groups. Based on this selection criteria, we also selected the putative oxidoreductase proteins Tmari 0118 and Tmari 0511 from our experiments, being a total of 12 proteins to be candidates of MSL synthases (Supplementary Table 17). Furthermore, we also propose that it is highly probable that the protein performing this reaction is also present in other MSL DA producers not related to Thermotogales (i.e., *Butyrivibrio fibrisolvens* or *Sarcina ventriculi*). Therefore, another criteria to select this group of proteins was based on the detection of homologous proteins in those MSL DA producers; in this regard, the above mentioned genes/proteins meet these criteria (Supplementary Figure 9, Supplementary Table 17).

To gain further insights and evaluate at the genomic level if the shared protein-coding genes among the reported DA producers code for proteins with a potential shared function (i.e. synthesis of DA), we identified the pangenome from a total of 54 strains, belonging to 10 species of identified DA producers belonging to Thermotogales, *Butyrivibrio* and *Sarcina*, and included one bacterial species *(E. coli*) not producing DA (Supplementary Table 18) as another selection criteria (i.e. Pangenome approach; Supplementary Figure 9). Based on this analysis, we evaluated if the upregulated protein-coding genes contained orthologous genes in most DA producers. The potential MSL DA synthases (Supplementary Table 17), the selection of which is based on the experimental evidence upon activation, the predicted biochemical activity, and presence of homologs in other MSL forming bacteria (*Butyrivibrio fibriosolvens* and *Sarcina ventriculi*) showed that Tmari 0511, Tmari 0827 and Tmari 1324 also possess orthologous genes in 35 strains belonging to 8 different species producing DA, but also in *E. coli.* The Tmari 0106, Tmari 0771, Tmari 0926, Tmari 1545, Tmari 0180 and Tmari 1609 were present in 34 strains belonging to at least eight different species producing DAs, and none of these harbor orthologous genes in *E. coli*. In addition, Tmari 0118, Tmari 1330 and Tmari 1331 only have orthologous genes in less than 21 strains belonging to 5 species from the Thermotogales. Genome resequencing of *T. maritima* MSB8 REF[24,25] has resulted in different annotations and codes for this organism (i.e, TMX, THEMA_X, Tmari_X, THMA_X). The genes and proteins identified here were mapped to the most recent reference genome[25] utilizing the Tmari codes.

Some other radical SAM proteins or oxidoreductases that were also upregulated in our experiments without a potential homolog in either one or both of the genomes of the DA producers *B.*

*fibrisolvens* or *C. ventriculi* were detected (i.e., Tmari 0114, Tmari 0422, Tmari 0825, Tmari 0950, Tmari 1290, Tmari 1296 and Tmari 1305 and Tmari 1347) (Supplementary Table 18). Other selection criteria for targeting the enzymatic process involved in the synthesis of MSLs, points to the lipid substrate (Supplementary Figure 9). Given that the biosynthetic precursors producing MSLs (i.e. FA tails) are known to be embedded into the cytoplasmic membrane forming the hydrophobic domain, it has been proposed that this enzymatic reaction could be partially associated or due to an integral membrane protein[21]. As seen with the biosynthesis of glycerolipids, most of the reactions involved in forming the core glycerol lipid are known to be catalyzed by integral or membrane-associated proteins[26]. Therefore, genes coding for proteins or the proteins predicted to be located in the membrane (as classified by the TMHMM algorithm) (Krogh *et al.*, 2001) have also been selected from our experiments (Supplementary Figure 9). Among these protein-coding genes, predicted integral components of the membrane (Tmari 0224, Tmari0309, Tmari 0488, Tmari 0513, Tmari 0559, Tmari 0560, Tmari 0840, Tmari 0978, Tmari1139, Tmari1257, Tmari1314, Tmari1409, Tmari 522 and Tmari1815), some of them also have homologs in the genomes of other MSL DA producers (Supplementary Table 18). We also applied the pangenome analysis to the genes and proteins that were categorized as suspected to be involved in the biosynthesis of MSL DAs (Supplementary Table 18), which their selection was based on their cellular localization, their predicted biochemical function, but that we did not always detect any potential homologs in both, or any other MSL DA producer outside from Thermotogales. From this analysis, only Tmari 0224 and Tmari 1012 contained orthologous genes in *E. coli.* The Tmari 0224, Tmari 0825, Tmari 0922, Tmari 1560, Tmari 1139, Tmari 0108, Tmari 0580, Tmari 0488, Tmari 0840, Tmari 1012, Tmari 0114, Tmari 0309, Tmari 0560 and Tmari 1305 contained orthologs at least in 20 strains from 5 different species producing DA. The rest of the selected genes were found between 7 and 16 strains belonging to 2 or 4 species producing DA (Supplementary Table 18). From our analysis, we have ranked those genes/proteins due to their predicted biochemical activity, the cellular localization and the presence of homologous proteins in other species producing DA with the pangenome analysis approach. In this sense, Supplementary Table 18 overviews the genes/proteins with higher to lower probabilities to also be part of the molecular mechanism of MSL synthase, mainly supported by the shared function of these genes among species.

Lastly, in the case that the targeted enzymatic reaction could be performed by proteins that are unique to *T. maritima* or closely related bacteria, and not present in other species producing MSL DA, or that their functions are not yet classified into any of the expected biochemical predictions, this could represent other criteria for recruiting potential proteins involved in this novel biosynthetic mechanism, from which we possess only experimental evidence of gene or protein activation, or that we ranked through the pangenome analysis (Supplementary Table 19). The major issue with these criteria is that overall, many proteins remain to be biochemically uncharacterized or to discover their function, and therefore this classification would not be enough as an ultimate selection criterion itself to consider for experimental validation with gene overexpression or gene deletion. Nevertheless, a list containing those proteins and predicted membrane proteins that were activated in our experiments and cannot be attributed to any other biosynthetic pathway or biological function is provided (Supplementary Table 19).

#### Confirmation of the activity of the potential MSL synthases and ether-forming enzymes

##### 1) *Confirmation of the membrane-spanning lipid synthases*

All candidate MSL synthase selected from the omic analyses of *T. maritima* and *T. ethanolicus* (Supplementary Tables 16, 17), were cloned in the pET29b expression vector and expressed in *E. coli* BL21 DE3 as a host, and incubated for 3 and 16 h after induction with IPTG both under aerobic and anaerobic conditions. From the chosen list of the MSL synthase potential candidates in *T. maritima* and *T. ethanolicus*, only the candidate MSL synthase of *T. ethanolicus* with accession number WP_129545148.1 resulted in the formation of diacids (coupling of two fatty acids) after 3 h postinduction, and exclusively in anaerobic conditions (Figure 2). Protein gels were run with the protein extracts of *E. coli* BL21 DE3 expressing the 'empty' pET29b expression vector and the WP_129545148.1 of *T. ethanolicus* expressed both under aerobic and anaerobic conditions as shown in Supplementary Figure 11. The negative control of the *E. coli* BL21 DE3 host strain transformed with the 'empty' pET29b expression vector (without MSL synthase genes) did not show peaks in the 'diacid region' of the chromatogram (Figure 2a). On the other hand, the *E. coli* BL21 + pET29b including the radical SAM protein of *T. ethanolicus* (WP_129545148.1) resulted in the biosynthesis of multiple diacids. The two most abundant diacids were identified as C33:1

and C34:2 compatible with the presence of 1 and 2 unsaturation, respectively. These diacids would be in agreement with a coupling of a C16:0 FA and a C17:1 FA and two times a C17:1 FA, respectively. To further investigate the nature of the unsaturations, the fraction containing the FAs and diacids (analyzed as their methyl esters) was hydrogenated. The C17:1 FA was unaffected by the hydrogenation, and its mass spectrum is compatible with the presence of a ring (Figure 2d). Comparing this data with that of REF[27], we concluded that the C17:1 FA has a cyclopropyl ring between position 9 and 10. This is indeed an abundant FA of *E. coli*, which was also detected in the control experiment. The C33:1 and C34:2 diacids also remained unaffected by the hydrogenation. Consequently, the two abundant diacids detected in the lipid extract of *E.coli* BL21 + pET29b_*T. ethanolicus*_WP_129545148.1 were identified as a C33 diacid containing one cyclopropyl ring (cyC33:0), and a C34 diacid containing two cyclopropyl rings (2x cyC34:0) (see Figure 2d). Other diacids were also detected in minor proportions including C30:0, C31:1, C32:0, C32:1, C33:1, C33:2, C35:1, C36:2 diacids (Supplementary Figure 12). Based on the mass spectral characteristics and relative retention time data of the C30:0 diacid this compound was identified as a diacid with a linear (i.e., unbranched) chain. Considering the FA composition of the *E. coli* BL21 expression host[28], the formation of the C30:0 diacid would be compatible by the coupling of a C14:0 FA and a C16:0 FA, generating a C-C bond between the ω positions of their alkyl chains. Similarly, the cyC33:0 and 2x cyC34:0 diacids must have been produced by ω-ω coupling of a n-C16 FA and a cyC17 FA and two cyC17 FAs, respectively.

It should be noted that the most closely related homolog to the MSL synthase of *T. ethanolicus* (WP_129545148.1) in the *T. maritima* genome does not correspond to any of the top hits included in Supplementary Table 17 that were tested by heterologous gene expression in *E. coli*, but rather to another hit (accession number WP_004080829.1, Tmari_0825, see Supplementary Table 18) also upregulated but not among the top MSL synthase candidates. It is important to highlight that this gene (Tmari _0825) is adjacent to the gene Tmari_0827 encoding the protein WP_004080828.1 (Supplementary Figure 13), which was among the top hits for potential MSL synthases (Supplementary Table 17). The protein coded by Tmari_0827 did not lead to the formation of diacids when expressed in *E. coli* BL21 DE3. Tmari_0825 was cloned in the expression vector pCDFDuet-1, transform it and express it in *E.coli* BL21 both under aerobic and anaerobic conditions, which lead to the synthesis of C32 DA and a cyC33 DA only under anaerobic conditions (Figure 2b, d). Hence, we conclude that this protein is responsible for the coupling of two FAs by ω-1/ω-1 of their alkyl chains.

Protein gels were run with the protein extracts of *E. coli* BL21 DE3 expressing the 'empty' pCDFDuet-1 expression vector and Tmari_0825 expressed both under aerobic and anaerobic conditions as shown in Supplementary Figure 11. In addition, due to the discrepancies between the different annotations of the genome of *T. maritima*, we have listed the correspondence of the accession number protein with previous annotations and other databases in Supplementary Table 20.

**2) *Confirmation of the ether-bond forming enzyme***

The modified-*plsA* gene of *T. maritima* (Tmari_0479), which we hypothesized to be involved in the formation of alkyl ether bonds of glycerol monoethers[21], and found to be activated by transcriptomics and proteomics in the current study (Supplementary Table 14), was also heterologously expressed in *E. coli* BL21 DE3. We also tested the expression of different predicted modified-*plsA* genes proposed in REF[21]: modified-*plsA* from *Desulfatibacillum alkenivorans*, *Thermodesulfobacterium geofontis* and *Candidatus* Kuenenia stuttgartensis (see Supplementary Table 21 for accession numbers). Expression of these modified-*plsA* coding genes resulted in the formation of *sn1*-alkyl glycerol monoethers in the case of *T. maritima* and *D. alkenivorans* only under anaerobic conditions (Figure 3, Supplementary Figure 14). In the case of *T. maritima*, only the C16:0 alkyl glycerol monoether was detected, while in the case of *D. alkenivorans* a C14:0, C16:0, cyC17:0 and C18:1 alkyl glycerol monoethers were formed (Figure 3, Supplementary Figure 14). Although both *Thermodesulfobacterium geofontis* and *Ca.* Kuenenia stuttgartensis modified-plsA proteins have a similar protein domain architecture of that found in *T. maritima* and *D. alkenivorans* (Supplementary Table 21), alkyl glycerol monoethers were not detected under the tested conditions. Protein gels of the cell extracts are shown in Supplementary Figure 15.

#### The widespread occurrence of MSL production in the Domain Bacteria

We screened selected genomes of bacteria with a confirmed presence/absence of DA or *iso*-DA for the presence of homologs of the confirmed MSL synthase. This group of bacteria has either been analyzed previously in our laboratory with methods enabling the detection of DAs or *iso*-

DAs or uses literature data generated by laboratories where such methods were also applied. Conventional FA analysis will usually miss the presence of the higher molecular weight diacids. We also analyzed various strains of the Clostridia class for this study. Of the 18 examined strains, *Sulfobacillus thermosulfidooxidans*, *Desulfospicio gibsoniae*, *Alkaliphilus transvaalensis*, and *Oscillibacter valericigenes* of the Eubacteriales, *Halocella cellulosilytica* of the Halanaerobiales, and *Fervidicola ferrireducens*, *Thermosediminibacter oceanii* of the Thermodesiminibacterales, and *Thermoanaerobacter wiegelii*, *Morella thermoacetica*, *Caldicellulosiruptor owensensis* and *Caldanaerobacter subterraneus* of the Thermoanaerobacterales, synthesize *iso*-FA in different relative abundances (Supplementary Table 22). A high percentage of *iso*-$C_{15}$ was particularly associated with the two species of the Eubacteriales and Thermosediminibacterales and four out of five species of the Thermoanaerobacterales. In case of *Caldicellulosiruptor owensensis*, also belonging to the Thermoanaerobacterales, the core lipids comprised only 5% *iso*-$C_{15}$, but 38% *iso*-$C_{17}$ (Supplementary Table 22). $C_{30:0,}$ $C_{32:0}$ and $C_{34:0}$ *iso*-DAs were detected (Supplementary Table 22). $C_{30}$ *iso*-DA was detected in the Thermosediminibacterales *Fervidicola ferrireducens* and the Thermoanaerobacterales *Thermoanaerobacter wiegelii, Thermoanaerobacter ethanolicus, Moorella thermoacetica, Caldanaerobacter subterraneus* but not in *C. owensensis.* Interestingly, *C. owensensis* only produced the $C_{34}$ *iso*-DA. Since its FA distribution is dominated by *iso*-$C_{17}$ FA rather than *iso*-$C_{15}$ FA, this is in strong agreement with the hypothesis that *iso*-diabolic acids are biosynthesized by the combination of two iso FAs. *C. subterraneus* also contained a low percentage of a $C_{32}$ *iso*-DA, and $C_{30}$, $C_{32}$ and $C_{34}$ α,ω-diols and also two ω-hydroxy fatty acids. Despite being a thermophilic Thermosediminibacterales species, with 36% *iso*-$C_{15}$ FA in its core lipids, *Thermosediminibacter oceani* did not produce any *iso*-DAs (Supplementary Table 22). For the first time the presence of DA in two species of the phylum Dictyoglomi was also detected.

To screen the selected genomes for the presence of homologs of the confirmed MSL synthase, we performed protein Position-Specific Iterative (PSI)-BLAST searches, and we considered as homologs those with an e-value =< $1e^{-50}$ and identity ≥ 30%. These are stringent criteria as radical SAM proteins are identified based on the sequence motif CxxxCxxC and less-stringent BLAST analyses could potentially lead to matches with radical SAM homologs not involved in the formation of MSLs. The screened strains included members of different subdivisions of the Acidobacteria phylum (where some subdivisions produce *iso*-DA and other subdivisions do not[29,30]), and cultures of the Clostridia class (Terrabacteria group, phylum Firmicutes) analyzed in the current study, and some others previously reported, i.e. *Thermoanaerobacter* species, *Butyrivibrio fibrisolvens* and *Sarcina (Clostridium) ventriculi*, all within the class Clostridia (Supplementary Table 21), as well as others species within the Proteobacteria, Chloroflexi Verrucomicrobia, FCB superphylum, Dictyoglomi, PVC group, Aquificae, and Fusobacteria (see main text for details).

An alignment of some of the identified homologs of the *T. ethanolicus* (WP_129545148.1) and *T. maritima* (WP_004080829.1) MSL synthases in MSL producers (*iso*-DA and DA), revealed six conserved blocks among all the radical SAM proteins (Figure 4a, Supplementary Figure 16). Block I containing the cysteine motif (CxxxCxxC) showed that in addition to the conserved cysteine signature, asparagine and glycine residues are also conserved among the radical SAM protein of MSL producers, leading to the CNxxCxGC motif (Supplementary Figure 16). Block II contains ca. 42 amino acids, a glycine-rich region (or GGE motif)[31] corresponding to the SAM binding motif. This region also possesses leucine, a conserved phenylalanine and asparagine. Block III comprises *ca.* 25 amino acids and contains conserved asparagine, serine, and another glycine-rich region including arginine. Block IV, contains conserved glycine, serine, asparagine, and glutamic acid. An aromatic tryptophan residue, followed by conserved proline, glutamine and arginine amino acids. Block V, possesses a conserved aspartic acid next to phenylalanine residues which are known to interact with FAs, as well as a tyrosine, histidine and asparagine surrounded by glycine moieties. A conserved VEPC motif a conserved proline, cysteine and phenylalanine at the end of the block. Finally, block VI contains three conserved proline residues and a cysteine, valine and alanine. The substrate-binding site in all radical enzymes suggests it is formed between the core region and structural domains localized mainly outside the core and near the C-terminus[31]. Hydrophobicity amino acid profile analysis[32] between the homologs of the MSL synthase showed enriched hydrophobic regions near the C-terminus of the proteins in all the *iso*-DA producers (Supplementary Figure 16). In contrast, the C-terminus of the radical SAM protein within the DA producers, showed less enriched hydrophobic regions (Supplementary Figure 16). The main differences between the radical proteins involved in the biosynthesis of MSL is likely related to the lipid substrate and the position of the carbon atom that is joined to another carbon atom of the lipid compound. In this sense, in the case of *iso*-DA producers the alkyl moiety is expected to derive from branched-chain FAs (i.e. *iso*-C15, *iso*-C17) joined at the ω carbon atoms, while in the DA producing MSL synthase the alkyl moieties are linear chains derived from *n*-FAs (i.e. C14,

C16), which are joined at the ω-1 carbon atoms. These differences between hydrophobicity of the radical SAM proteins in *iso*-DA and DA producers may be necessary to accommodate and bind the different lipid substrates and allow the hydrogen abstraction during the catalysis. Furthermore, it could be speculated that the carbon position at which the carbon-carbon bond is catalyzed might be determined by this region (Figure 4b, Supplementary Figure 16).

Structure-based sequence alignments indicate that the MSL synthase radical proteins possess the conserved β/α topology (Figure 4a, Supplementary Figure 16). A three-dimensional model prediction of the structure of the MSL synthases of *T. ethanolicus*, representing the *iso*-DA producers, and of *T. maritima*, representing the DA producers (Figure 4b, Supplementary Figure 17, a,b) showed what seems to be the common fold for the radical SAM proteins; the β/α motifs are arranged as a TIM-barrel[31]. In both cases, the radical SAM core is located near the N-terminus of the molecule with the conserved cysteine motif and the SAM binding site on a lateral opening of the barrel (Block I and Block II; Supplementary Figure 17a, lower panel). The C-terminal region of the radical SAM proteins are known to contribute to substrate specificity[33], in order to accommodate the substrate to the binding (active) site, the conformation and the substrate position is likely to be a prerequisite to ensure the proper hydrogen-atom abstraction during the reaction. In this sense, we have detected enriched hydrophobic regions within the N-terminal end of the proteins (Supplementary Figure 16). Structural conformation of both MSL synthases indicates this region localizes near the radical SAM active site, suggesting that this portion of the protein chain? (Supplementary Figure 16b,c, Supplementary Figure 17a,b left lower panel) is likely to interact with the methyl-branched FAs (*iso*-C15) in the case of iso-DA producers and with straight chain FAs (C16) in the case of DA producers. However, it is possible that conformational changes could occur upon substrate binding as seen with other radical SAM enzymes[31] and that additional regions may interact with both tails of the corresponding FAs.

We extended the screening of MSL synthase homologs in bacterial genomes compiled in the NCBI non-redundant protein sequence database using the same stringent blast criteria and built a maximum likelihood phylogenetic tree (Figure 5a, Supplementary File 1). Homologs were detected in species of the phyla Firmicutes (classes Clostridia, Tissierellia, Erysipelotrichia, Bacilli, Negativicutes, Limnochordia), Actinobacteria (class Coriobacteriia), Thermotogae, Synergistetes, Spirochaetes, Armatimonadetes, Caldiserica, Calditrichaeota,

Coprothermobacterota, Chloroflexi, Nitrospirae, Elusimicrobia, Deferribacteres, Dictyoglomi, Proteobacteria (gamma and delta), Atribacterota, FCB superphylum (Bacteroidetes, Ignavibacteriae, Fibrobacteres, Candidatus Zixibacteria, Latescibacteria, Marinimicrobia, Cloacimonetes), Cyanobacteria/Melainabacteria group, PVC group (phyla Omnitrophica, Planctomycetes), and in multiple members of the bacteria candidate phyla. The MSL homologs within the phylum Firmicutes Clostridia class, were detected in the orders Eubacteriales, Thermoanaerobacterales, Thermosediminibacterales, Halanaerobiales of the class Clostridia, which suggests that other classes within the Clostridia class not screened for their membrane lipid composition might also harbor the capacity to biosynthesize MSLs.

It is important to note that despite of the apparent divergence of potential MSL producers in different phylogenetic groups, based on the presence of the MSL synthase homolog (Supplementary Table 21, Figure 5a, Supplementary File 1), the phylogeny of Clostridia, among others groups mentioned here, is still quite unresolved and most of these potential MSL producers are somewhat related. For one, despite being Dictyoglomi assigned to their own phylum, some studies based on different genomic analyses methods[34,35] have reported that members of this phylum are closely related to Thermotogales and to the Clostridia Thermoanaerobacterales, and that might form a monophyletic group with Coprothermobacter, where we have also detected MSL synthase homologs (Figure 5a, Supplementary File 1). The same applies to the phylogenetic positioning of Syntrophomonadaceae and Halanaerobacteriales within the Clostridia, and the phylum Synergistetes[36], making it difficult to infer where and how the MSL synthesis capacity was acquired and potentially transferred to other phyla.

The detection of homologs of the MSL synthase in genomes of the *Candidatus* Cloacimonetes of the FCB superphylum (Figure 5a) is also remarkable as our previous study[37] detected and validated the activity of homologs of the archaeal membrane lipid biosynthesis in members of *Ca.* Cloacimonetes, which also harbored the canonical bacterial FA biosynthetic pathway. The presence of the bacterial MSL synthase homolog in *Ca.* Cloacimonetes genomes suggest the possibility that members of this group could also synthesize bacterial MSLs. MSL synthase homologs were also found in archaeal genomes, i.e, specific euryarchaeota groups (see main text for details), in (meta)genomes of members of the DPANN Archaea (*Ca.* Woesearchaeota, Aenigmarchaeota, Pacearchaeota), and members of the Asgard group (*Ca.* Lokiarchaeota, *Ca.*

Heimdallarchaeota, *Ca.* Thorarchaeota) (Figure 5a, Supplementary File 1). The presence of these homologs is enigmatic and further research is required to confirm their activity in Archaea (see main text for further discussion).

#### The widespread occurrence of ether lipid production in the Domain Bacteria

The capacity to synthesize bacterial alkyl glycerol ether lipids was also screened for in the list of selected bacteria for this study (Supplementary Table 21). The genomes of those strains were also screened for the presence of the homolog of the glycerol ester reductase (GeR) of *T. maritima* with the same astringent criteria used for the MSL synthase homolog search. We also investigated the functional domain architectures for those screened strains in the genomes of which we could detect a homolog to the confirmed GeR of *T. maritima*. Visual presentations of the different architectures are included in Supplementary Figure 18 and discussed in the main text. We also detected the GeR homolog in two Acidobacteria genomes, *Edaphobacter aggregans* (SD1, REF[30]) and in *Holophaga foetida* (SD8), both reported to synthesize glycerol ethers (Supplementary Table 21). Of all the Acidobacteria strains known to synthesize ether-bonded lipids, *E. aggregans* and *H. foetida* are either facultative anaerobes or strict anaerobes (together with *Thermotomaculum hydrothermale*, in the genome of which we detect the GeR homolog but has not been reported to synthesize alkyl ether lipids in culture) (Supplementary Table 21) (REF[38]). Therefore, we speculate that facultative or strict anaerobic groups within the Acidobacteria could potentially be producers of (MSL) ether lipids by using the GeR confirmed in this study. In order to further confirm this hypothesis, we performed PSI-pblast searches with the GeR of *T. maritima* (AGL49404.1) as a query against the non-redundant protein database by restricting to sequences affiliated to the Acidobacteria. Potential homologs were detected in acidobacterial genomes amplified from wetland soil, freshwater sediment, Siberian permafrost, and in peatland acidobacteria metagenome-assembled genomes of the *Ca.* Sulfotelmatomonas gaucii and *Ca. Sulfopaludibacter*[39]. This finding further supports the potential synthesis of ether-bonds by (facultative) anaerobic acidobacteria by using GeR.

We also investigated the presence of homologs of the GeR of *T. maritima* in bacterial genomes compiled in the non-redundant protein sequence database and we detected them in species of different bacterial and archaeal phyla as indicated in the main text (Figure 5b, Supplementary File

2). Among bacterial genomes, GeR homologs were detected in the phyla Firmicutes, Actinobacteria, Thermotogae, Synergistetes, Spirochaetes, Armatimonadetes, Calditrichaeota, Chloroflexi, Nitrospirae, Elusimicrobia, Deferribacteres, Proteobacteria (alpha, gamma and delta), Atribacterota, FCB superphylum (Bacteroidetes, Ignavibacteriae, Fibrobacteres, Candidatus Zixibacteria, Latescibacteria, Chlorobi, Marinimicrobia, Cloacimonetes, Gemmatimonadetes, Fermentibacteria), Cyanobacteria/Melainabacteria group, PVC group (phyla Omnitrophica, Verrucomicrobia, Planctomycetes), Nitrospinaea, Thermodesulfobacteria, Aquificae, Acidobacteria, Chryosiogenetes, and in multiple members of the bacteria candidate phyla (Figure 5b, Supplementary File 2).

#### Supplementary Figures

**Supplementary Figure 1.** Structures of (1) a branched glycerol dialkyl glycerol tetraether (brGDGT), (2) $C_{30}$ *iso*-diabolic acid (*iso*-DA) (13,16-dimethyloctacosanedioic acid) (3) the glycerol ether derivative of *iso*-diabolic acid and (4) $C_{32}$ diabolic acid (DA) (15,16-dimethyltriacontanedioic acid).

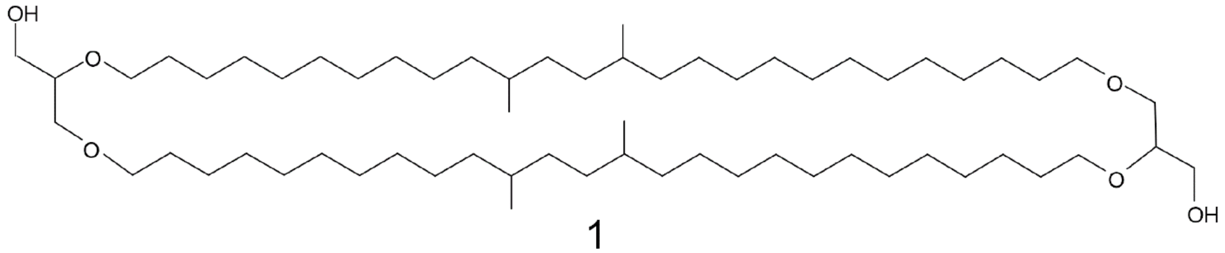


1

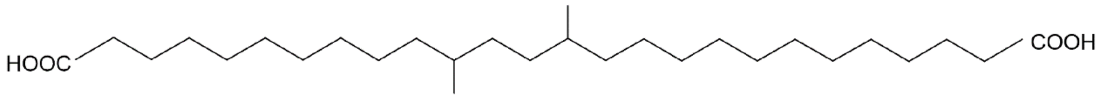


2

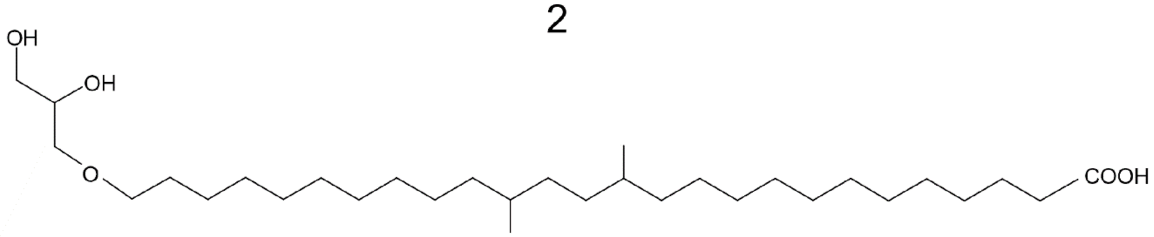


3

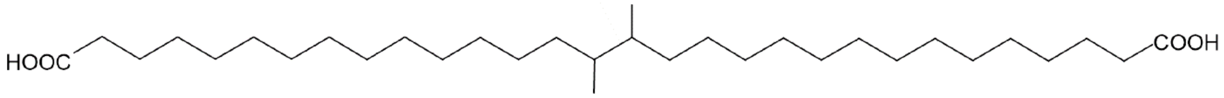


4

**Supplementary Figure 2.** Core lipid composition of *Thermotoga maritima* at optimal (80°C) or suboptimal (55°C) growth temperature during various growth phases. a) Core lipid distribution of C14, C16 and C18 fatty acids (FA) C30 and C32 diabolic acids (DA), fatty acids ether C16, 1-*O*-hexadecyl glycerol (GE), diabolic acids glycerol ethers C30 and C32 (DA-GE), monomethyl diacids (MM) C30 and C32 and monomethyl diacids glycerol ether C30 and C32 (MM-GE). Bars represent the mean fractional abundance. The error bars indicate the standard deviation of three biological replicates. Significant (ANOVA) are indicated ($p<0.05$). b) Molar mass changes (%) of total ester and ether lipids across growth phases. c) Molar mass changes (%) of total membrane-spanning and non-spanning lipids across growth phases. The error bar indicates the standard deviation of three biological replicates. Statistically significant differences from ANOVA are indicated ($p<0.05$) Molar mass changes are calculated on the flame ionization detector response to the number of carbon atoms of the derivatized compounds (methyl ester and trimethylsilyl groups).

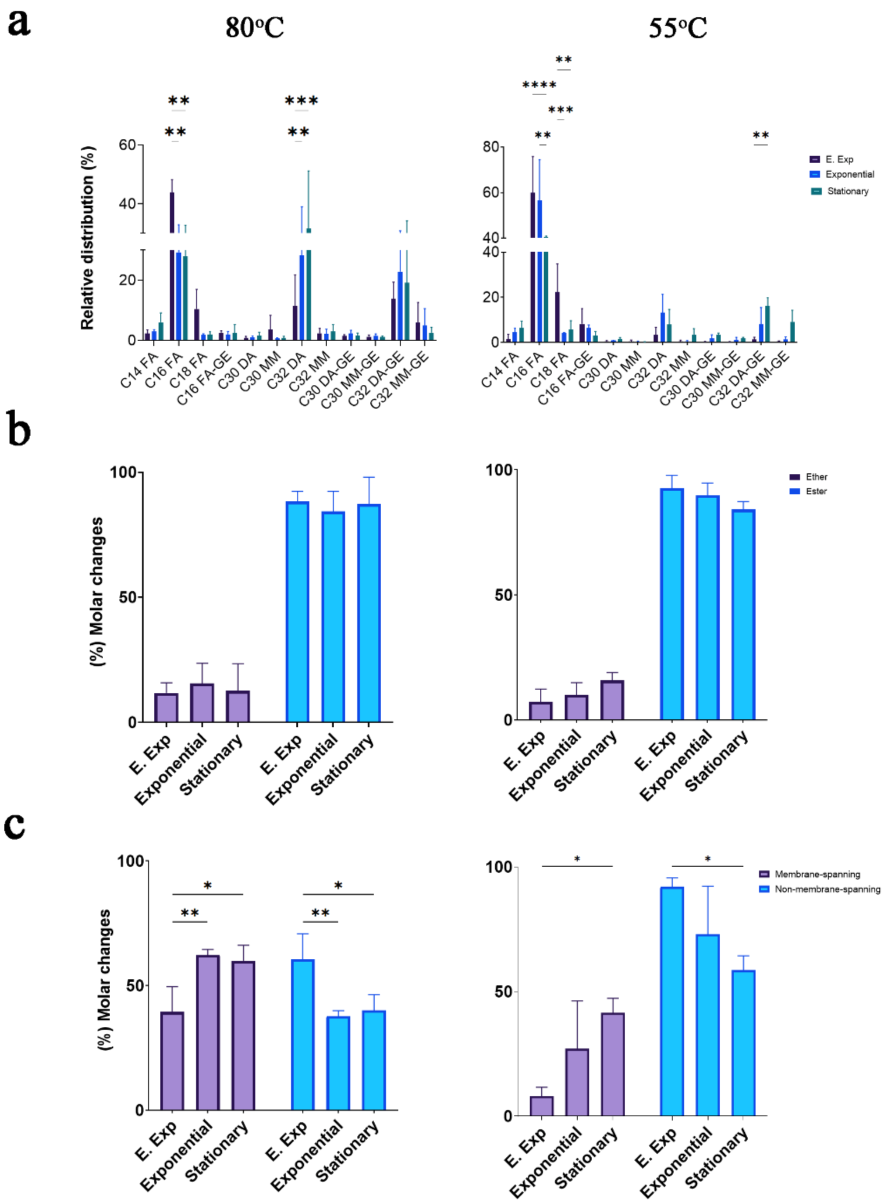

**Supplementary Figure 3.** Comparison of gene transcript abundances in *Thermoanaerobacter ethanolicus* cultures across growth phases at optimal 65°C and suboptimal temperature 45°C, by using Voronoi treemaps of the transcriptome data. Voronoi treemaps depict all significant transcripts giving the fold change according to the growth phase comparison. Big tiles symbolize major KEGG (Kyoto Encyclopedia of Genes and Genomes) classification: (front) small tiles represent single proteins (bottom). Expression is colored according to the abundance (red indicates more abundant, and blue is less abundant, grey, not detected) Growth phase comparisons are indicated.

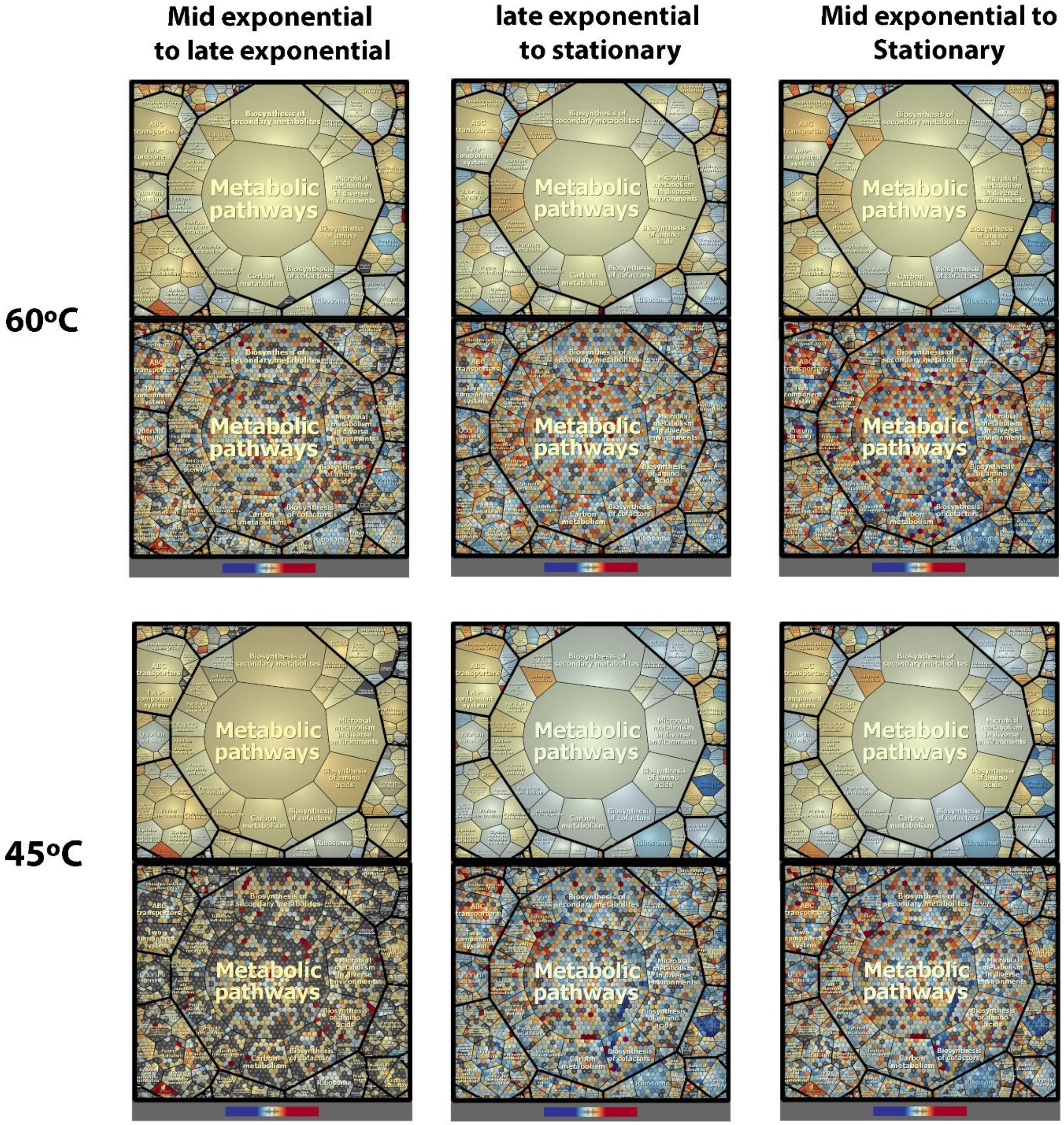

**Supplementary Figure 4.** Comparison of gene transcript abundances in *Thermoanaerobacter ethanolicus* cultures (transcriptome data) by taking into account the effect of temperature (60°C optimal *vs* 45°C suboptimal) displayed in Voronoi treemaps. Expression is colored according to the abundance (red indicates more abundant, and blue is less abundant, grey, not detected) Growth phases are indicated.

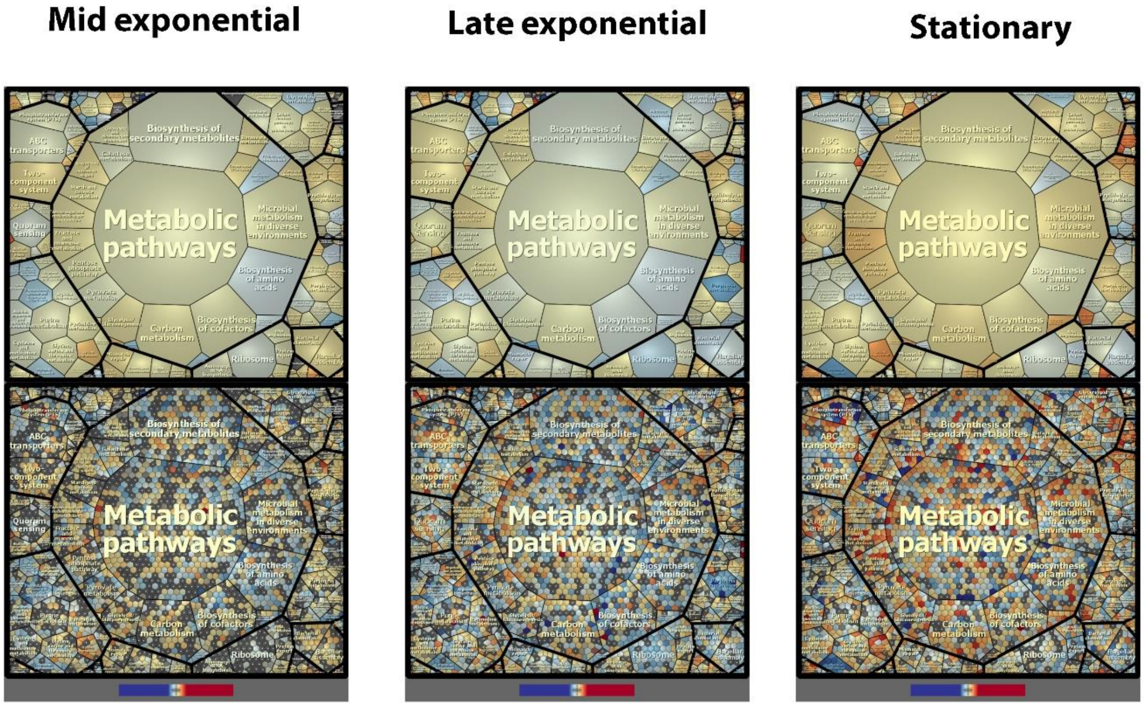

**Supplementary Figure 5.** Predicted biosynthetic pathway for lipid biosynthesis in *Thermoanaerobacter ethanolicus* based on the detected lipids and upregulation of genes in the transcriptomic data. Figure created using BioRender.

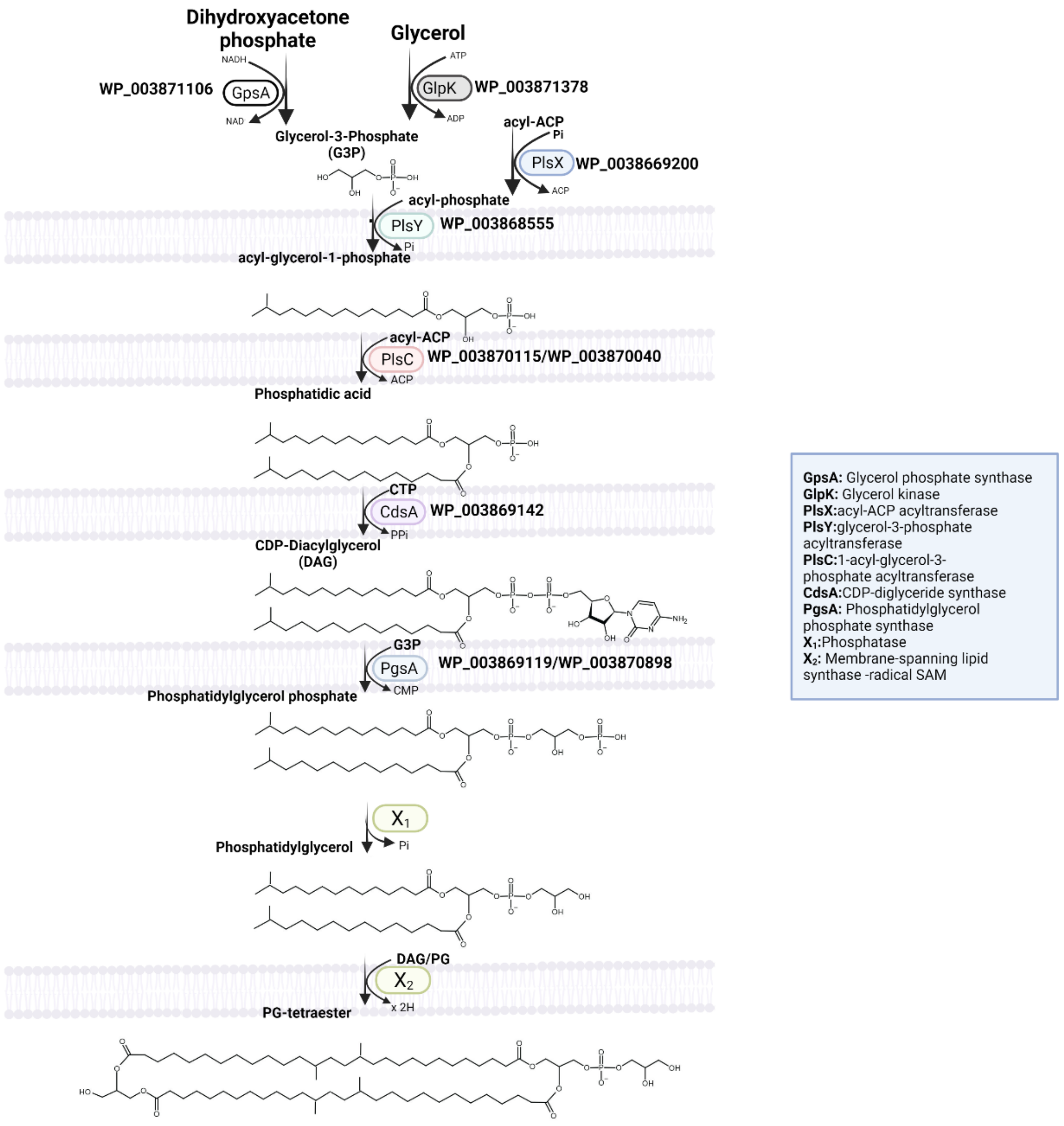

**Supplementary Figure 6**. Comparison of gene transcript and protein abundances in *Thermotoga maritima* cultures across growth phases at a) optimal 80°C and b) suboptimal temperature 55°C, by using Voronoi treemap of the transcriptome and proteome data. Voronoi treemaps depict all significant transcripts giving the fold change according to the growth phase comparison and of the detected proteins giving the protein ratio according to the growth phase comparison. Expression is colored according to the abundance (red indicates more abundant, and blue less abundant, gray, not detected). Growth phases comparisons and transcriptomic or proteomic analysis are indicated.

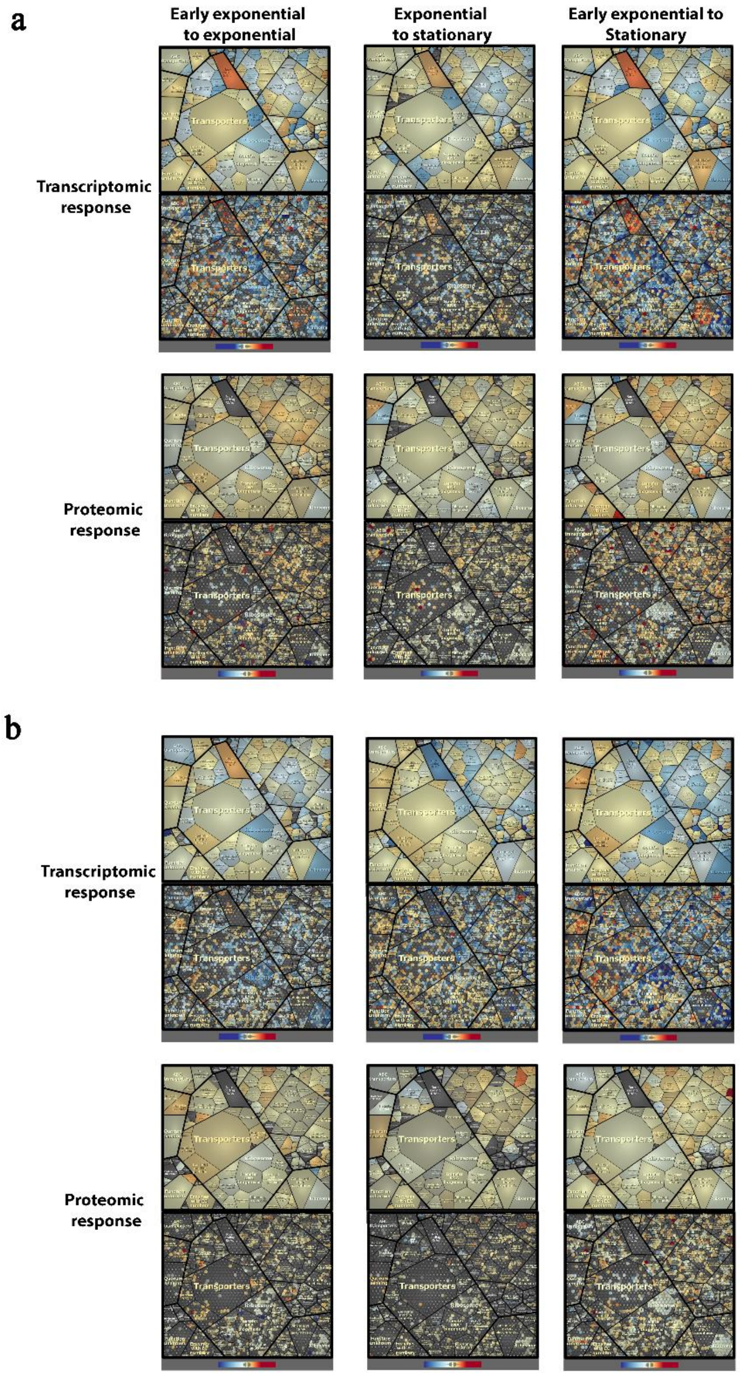

**Supplementary Figure 7.** Comparison of gene transcript and protein abundances in *Thermotoga maritima* cultures (transcriptome and proteome data) by taking into account the effect of temperature (80°C optimal *vs* 55°C suboptimal). Expression is colored according to the abundance (red indicates more abundant, and blue less abundant, gray, not detected). Growth phases and transcriptomic or proteomic analysis are indicated.

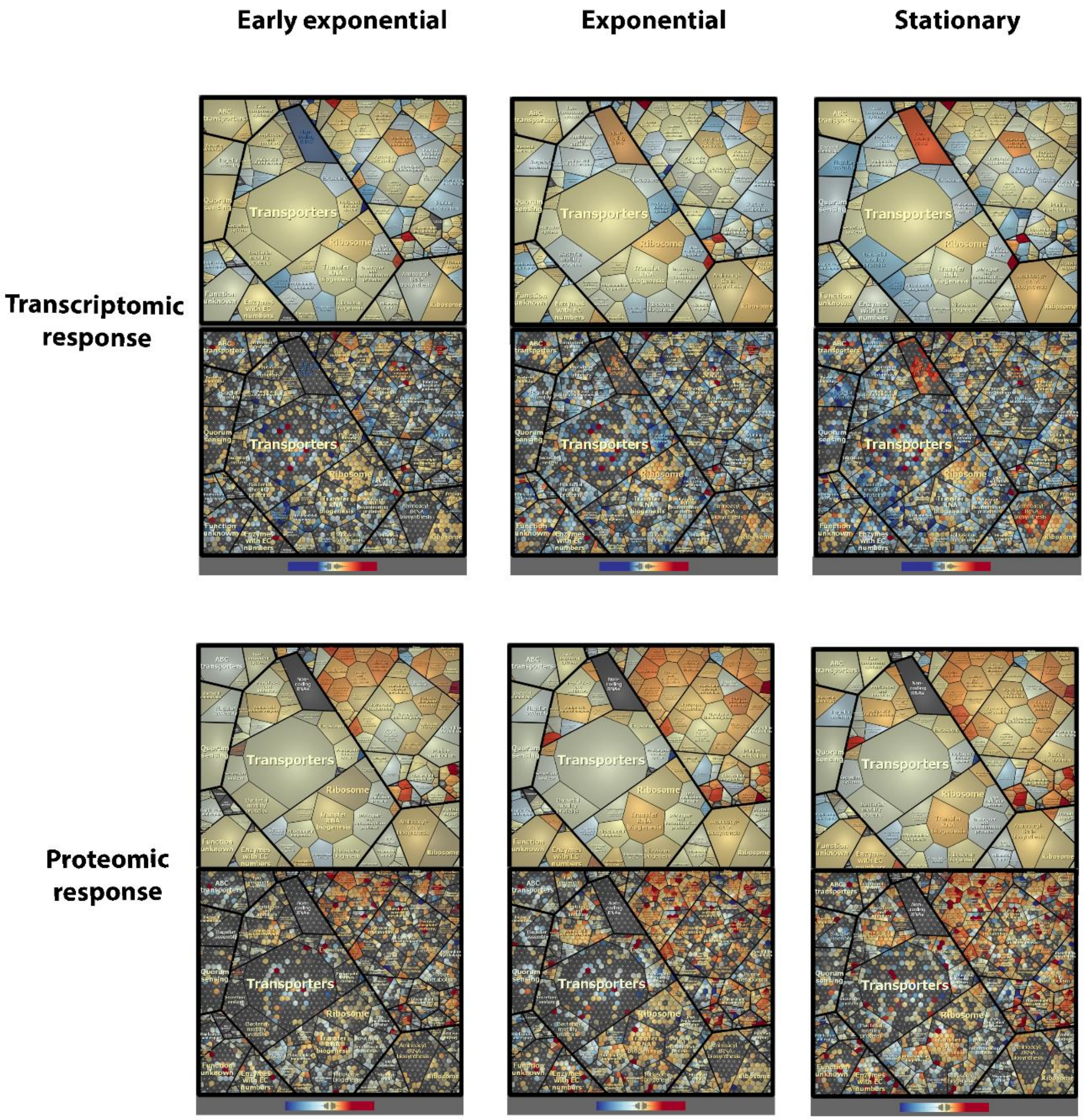

**Supplementary Figure 8.** Predicted biosynthetic pathway for lipid biosynthesis in *Thermotoga maritima* based on the detected lipids and upregulation of genes in the transcriptomic and proteomic data. Figure created using BioRender.

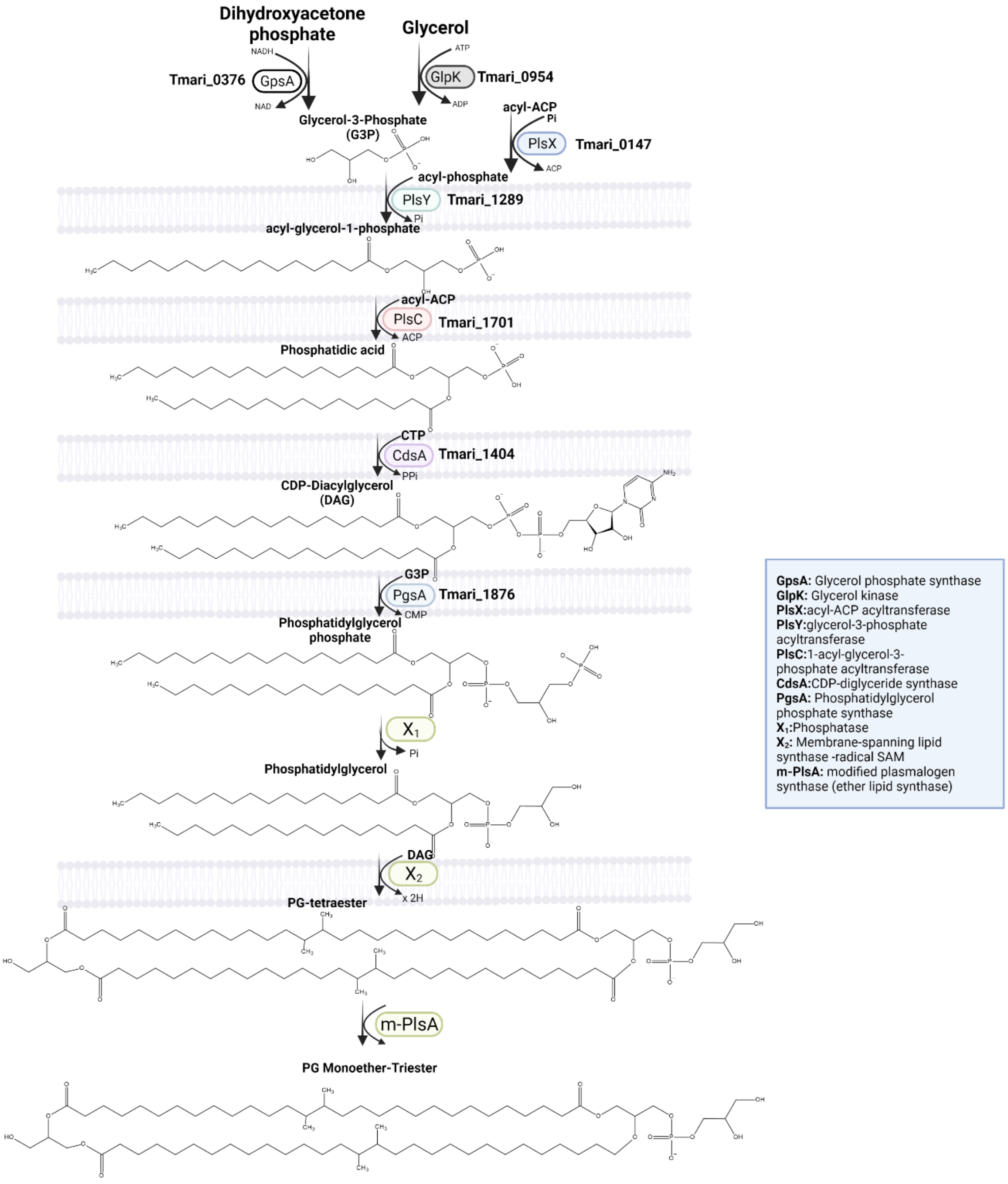

**Supplementary Figure 9.** Workflow for recruitment of potential MSL synthases in *Thermotoga maritima* and *Thermoanaerobacter ethanolicus* by using transcriptomic, proteomic, the pangenome approach, and selection criteria. Figure created using BioRender.

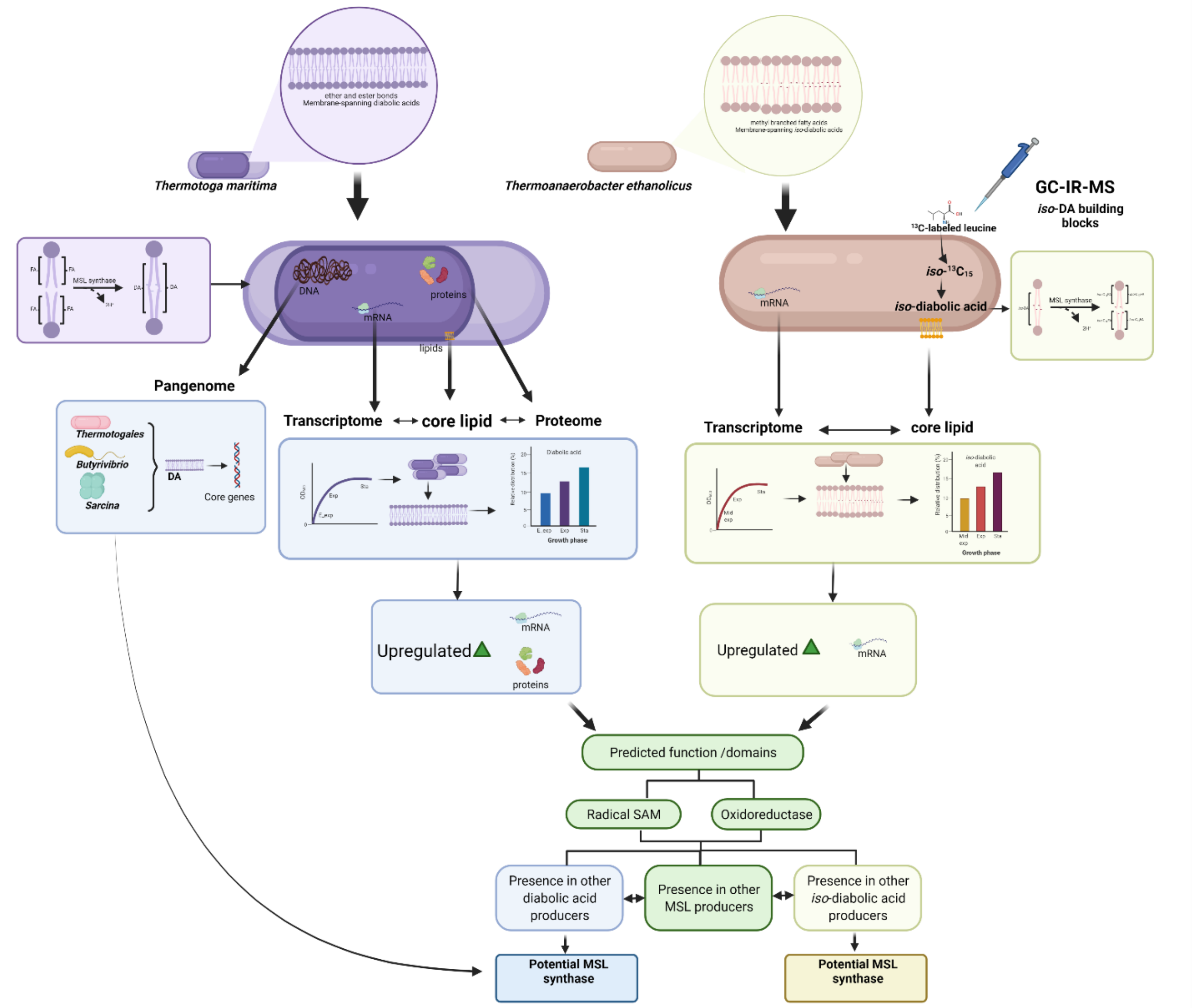

**Supplementary Figure 10**. Comparison of the *Thermoanaerobacter ethanolicus* MSL synthase WP_129545148.1 annotated protein in different annotation versions of the genome (see text for details). Amino acid position of the predicted functional domains are indicated above each box (showing the Pfam domain). Figure created using BioRender.

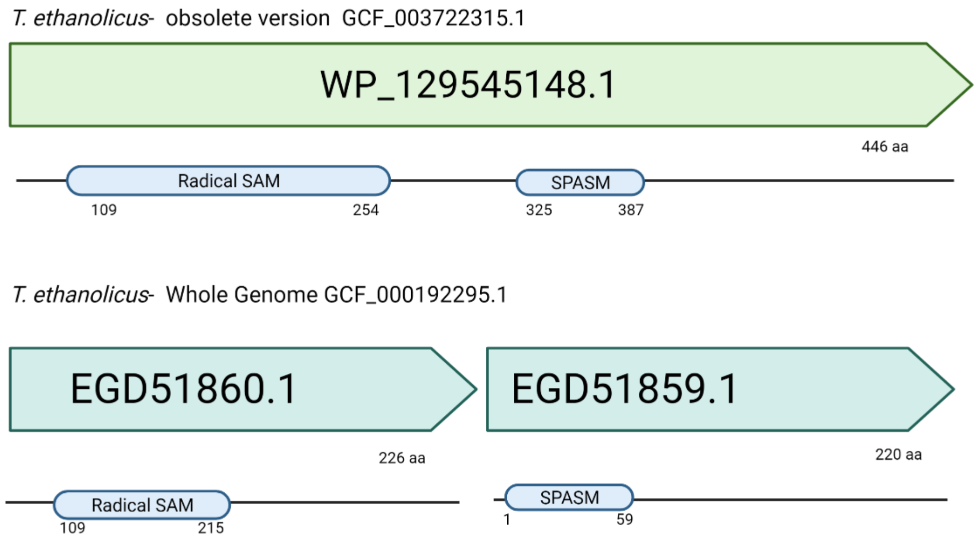

**Supplementary Figure 11.** SDS-PAGE gel images of the induced gene expression of the MSL synthase of *T. ethanolicus* (WP_129545148.1) in *E. coli* BL21 under anaerobic conditions, compared to the 'empty' vector pET29b, and cell extracts of *Thermoanaerobacter ethanolicus*. Lane 1, Molecular weight marker (10 to 250 kDaltons).

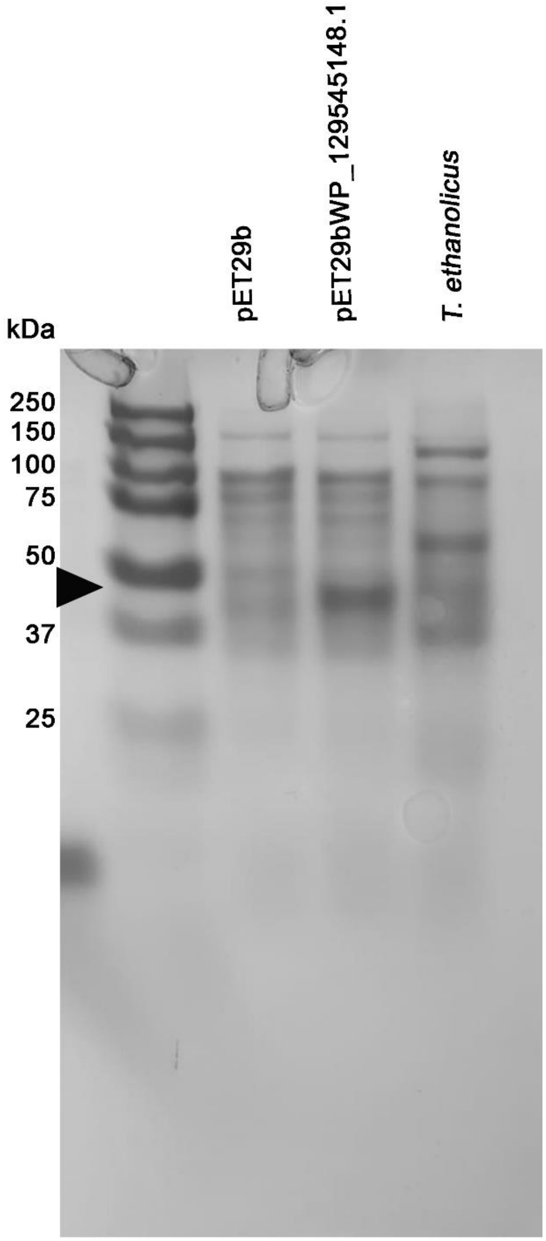

**Supplementary Figure 12.** Mass spectra (subtracted for background) of the dimethyl esters of eight diacids (in addition to those shown in Fig. 2a) detected in the *E.coli* BL21 upon expression of pET29b_WP_129545148.1 of *Thermoanaerobacter ethanolicus*.

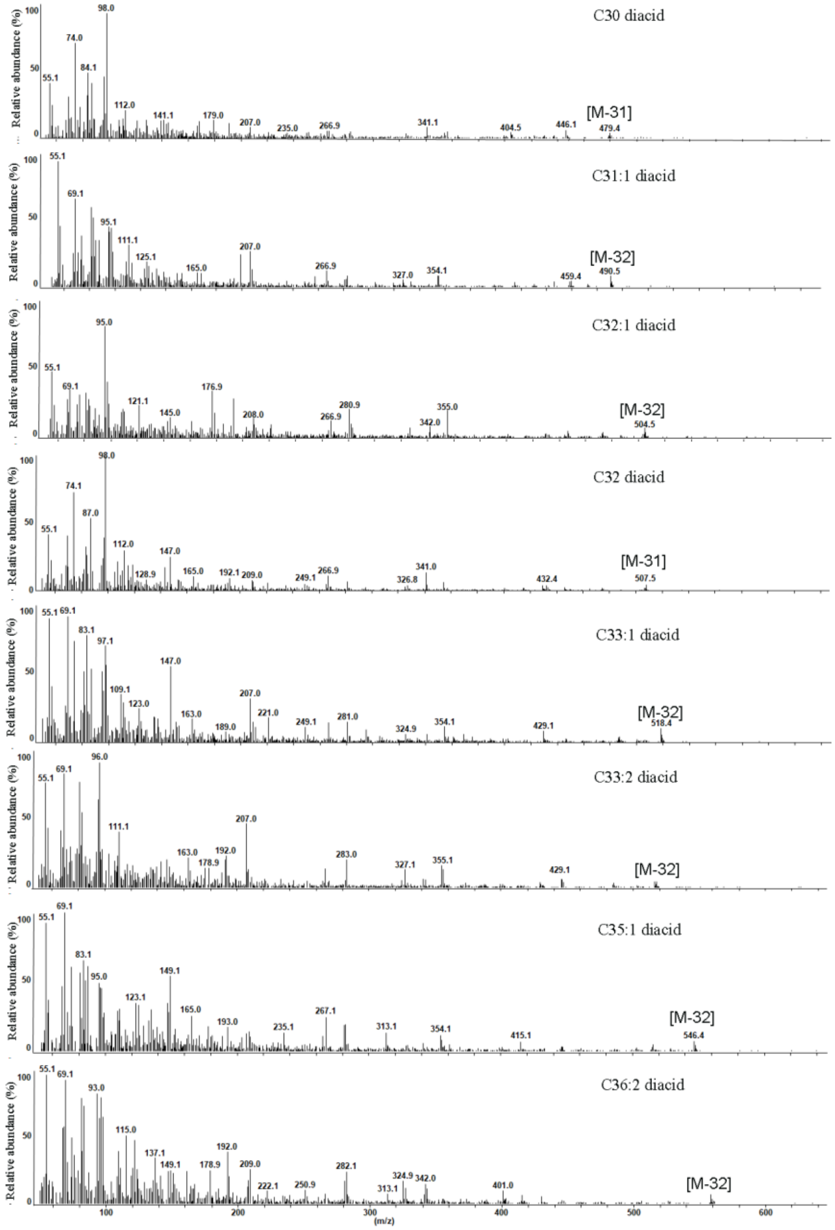

**Supplementary Figure 13.** Genomic localization of the potential MSL homolog of *T. maritima* (Tmari_0825) and in other members from the Thermotogales. Note the different annotation names are included (see Supplementary Table 20 for a detailed overview). The predicted domains are shown below the operon structure. Figure created using BioRender.

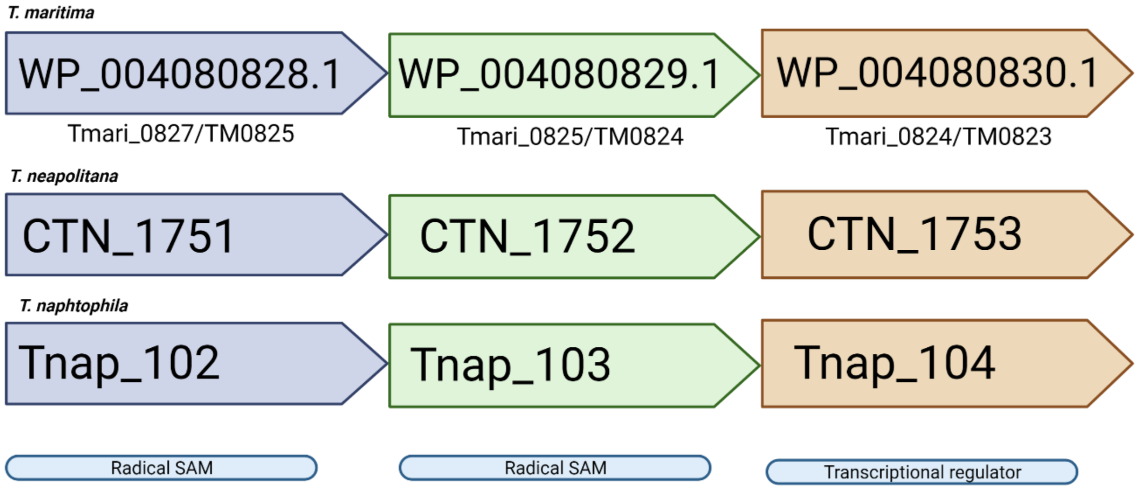

**Supplementary Figure 14.** Mass spectra of three additional 1-monoglycerylethers (1-MGE) with C14, cyC17 and C18:1 alkyl chains detected in the *E. coli* BL21 DE3 upon expression of the modified *pls*A gene (SHJ90043.1) (GeR) of *Desulfatibacillum alkenivorans.*

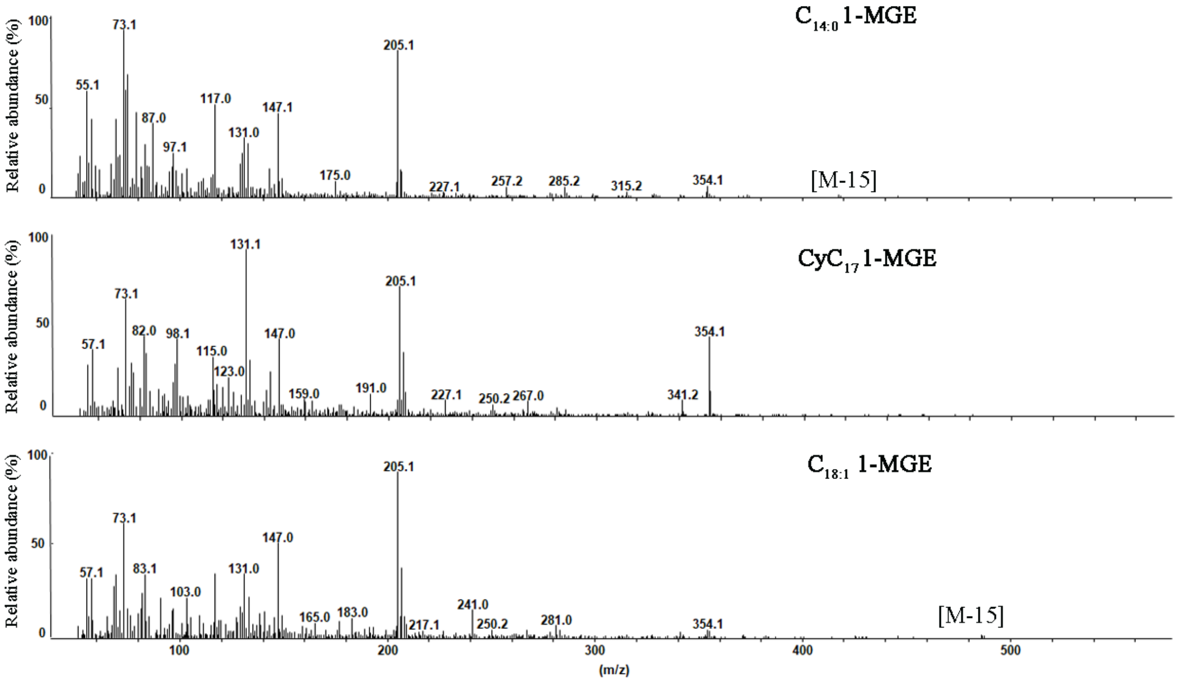

**Supplementary Figure 15**. SDS-PAGE gel images of the induced gene expression of the GeR of of *Desulfatibacillum alkenivorans* (SHJ90043.1), *Thermotoga maritima* (Tmari 0479), and of *Enterococcus faecalis* (AAO8118/EF_1327) in *E. coli* BL21 under anaerobic conditions after 3 or 14 hours postinduction. First lane, molecular marker indicating line of approx. 150kDaltons, size of expected protein.

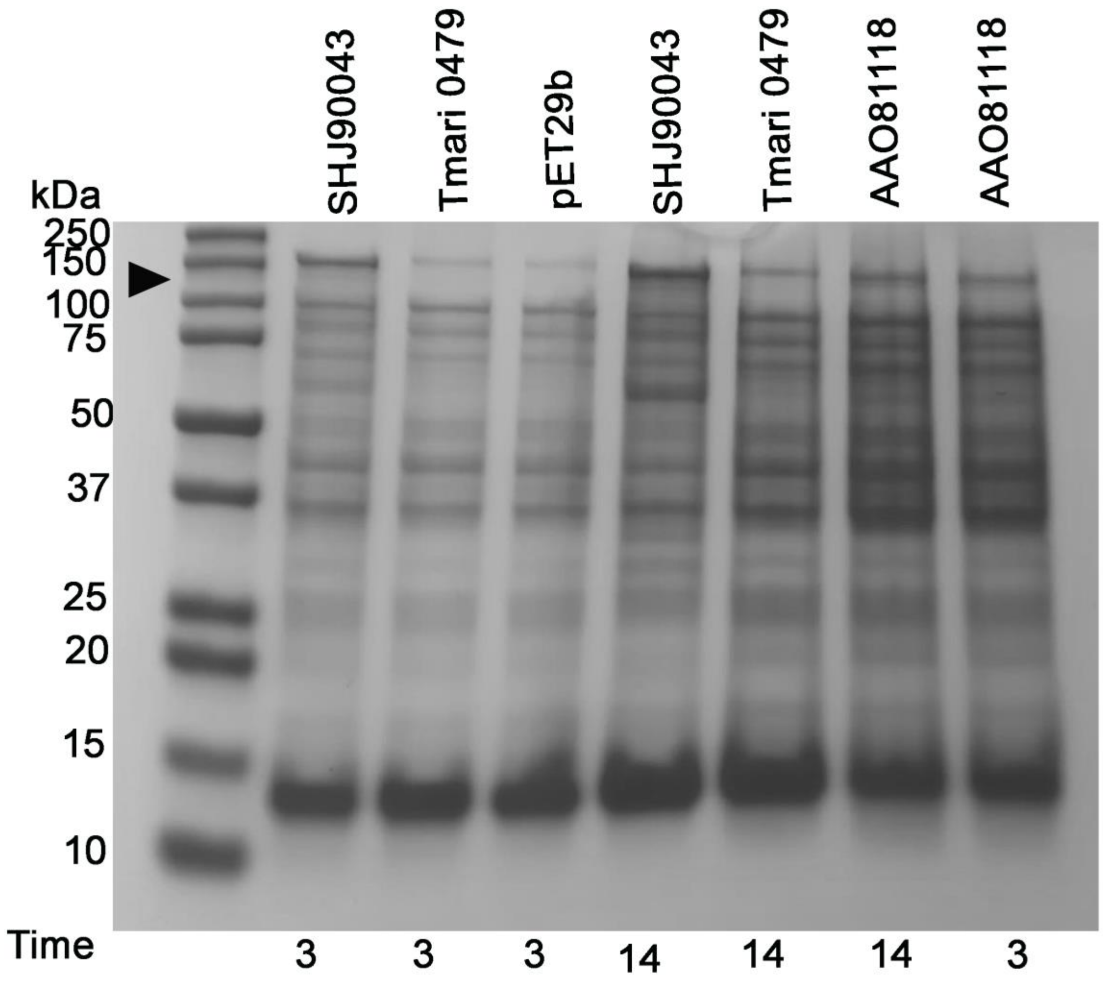

**Supplementary Figure 16.** Sequence alignment of the MSL synthases of *iso*-DA and DA producers. a) Conserved amino acid blocks of *Thermoanaerobacter ethanolicus* WP_129545148.1 and homologs of this protein (see Supplementary Table 21 for accession numbers) in other MSL producers. Amino acid block I possess the CxxxCxxC cysteine motif of the radical SAM proteins. The GGEP motif potentially involved in SAM binding is located in Block II together with a conserved leucine, glycine and a conserved phenylalanine. Block III possesses a conserved serine, arginine and asparagine and three conserved glycines along the block. Block IV comprises glycine, serine, asparagine, glutamate and tryptophan. Block V harbors a conserved VEPC motif. Block VI comprises three conserved proline, cysteine and valine. Identical residues defining the six blocks are shaded in blue. Structure-based sequence alignment of b) *iso*-DA and c) DA homologs protein to WP_129545148.1 of *T. ethanolicus.* In red hydrophobic residues are highlighted, and in blue the hydrophilic residues conserved among each MSL group. Predicted secondary structure for *T. ethanolicus mslS* (b) and for *T. maritima mslS* (C) is shown above the alignment. JNetPRED consensus prediction; helices are marked as red tubes, and sheets as green arrows.

**Supplementary Figure 16. Cont.**

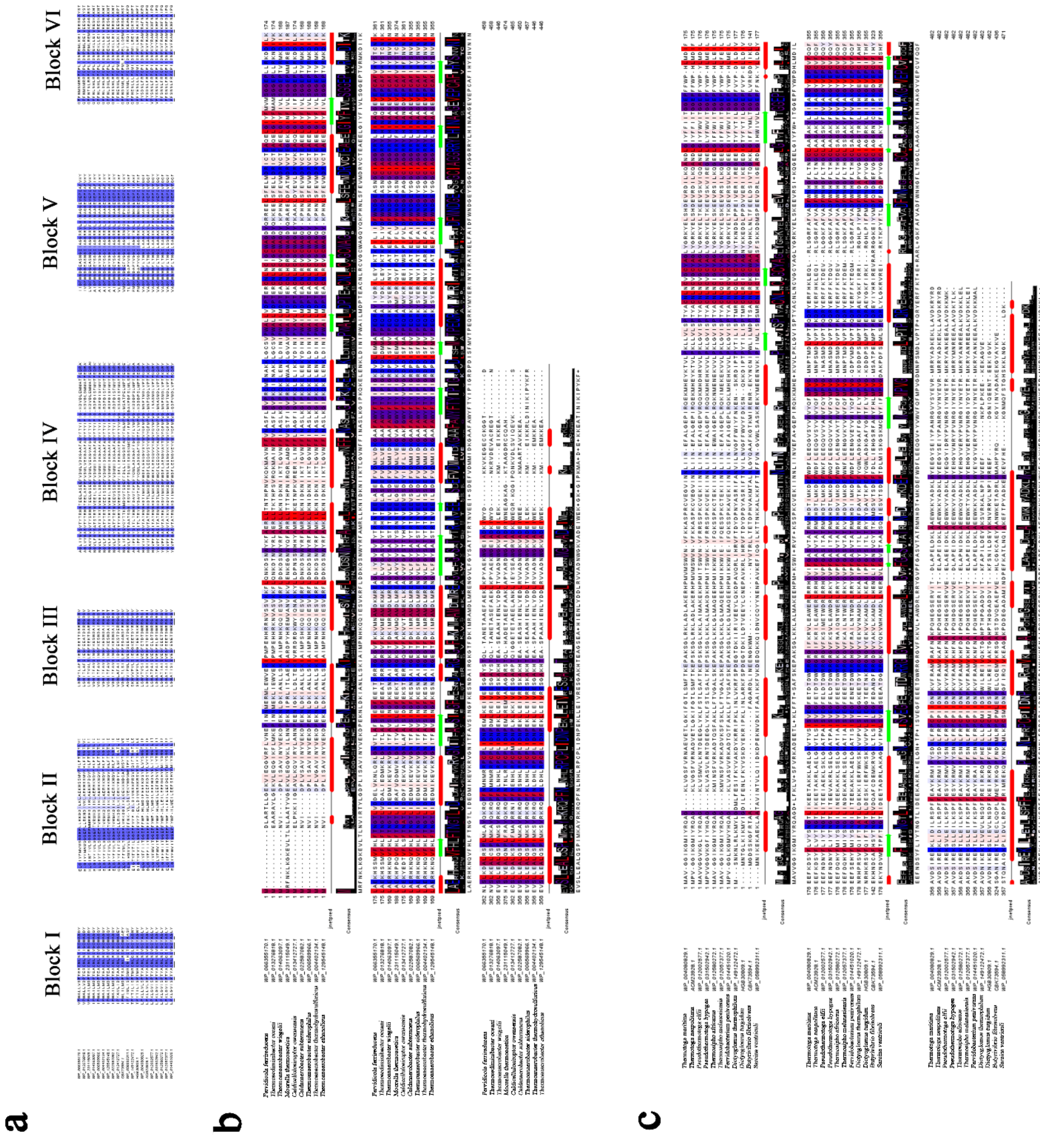

**Supplementary Figure 17.** 3D models of MSL synthases (MSLs) form a) *Thermoanaerobacter ethanolicus* and b) *Thermotoga maritima*. Representation of the predicted MSLs synthase tridimensional structure with the conserved cysteine motif involved in the iron-sulfur cluster (red) and the predicted SAM binding domain (blue) are represented as sticks. The protein (top) is represented in color gradient from N-terminal (blue) to C-terminal (red). Secondary structure is colored in the protein (bottom) The hydrophobic region potentially involved in substrate binding is colored in yellow and zoomed in.

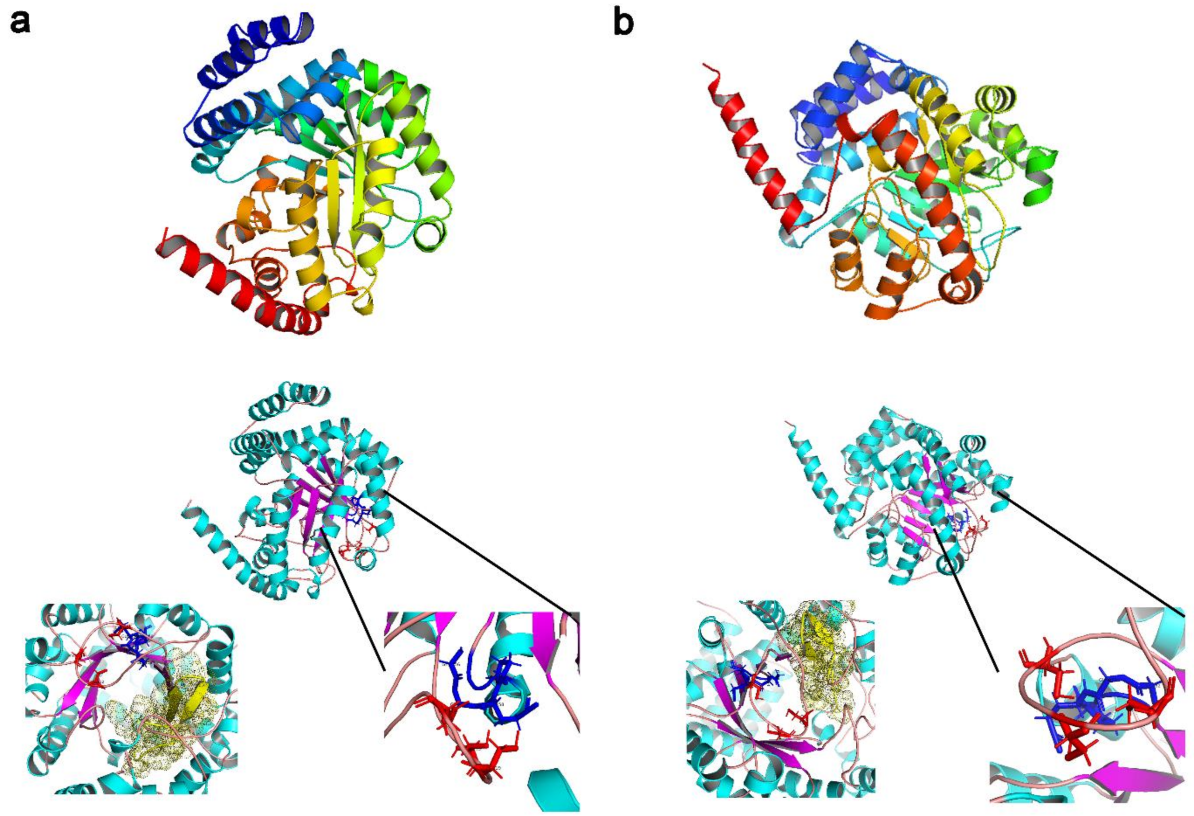

**Supplementary Figure 18.** Predicted architecture domain structure of the protein homologs to the glycerol ether reductase (GeR) of *T. maritima*. GeR was firstly identified as a modified-PlsA (plasmalogen synthase) protein homolog to the PlsA from *Enterococcus faecalis*, which is known to synthesize plasmalogens. *E. faecalis* architecture is composed of two activation domains (Benzoyl-CoA reductase PF01869+PF01869; *in red*) and two-electron transfer domains ( CoA enzyme activase uncharacterized domain PF09989; in yellow + 2-hydroxyglutaryl-CoA dehydratase, D component F06050; in blue, or the domain architecture 1. *T. maritima* GeR known to produce ether lipids (See Figure 3) contains architecture 1c along the protein. *Desulfatibacillum alkenivorans* GeR, also producing ether lipids (See Figure 3, Supplementary Figure 14) contains architecture 1i along the protein. Bacterial species with ether bond forming capacity contains different domain organization along the GeR protein homolog (See Supplementary Table 21). Analysis performed with Interpro[40] scan and Pfam[41]. Figure created using BioRender

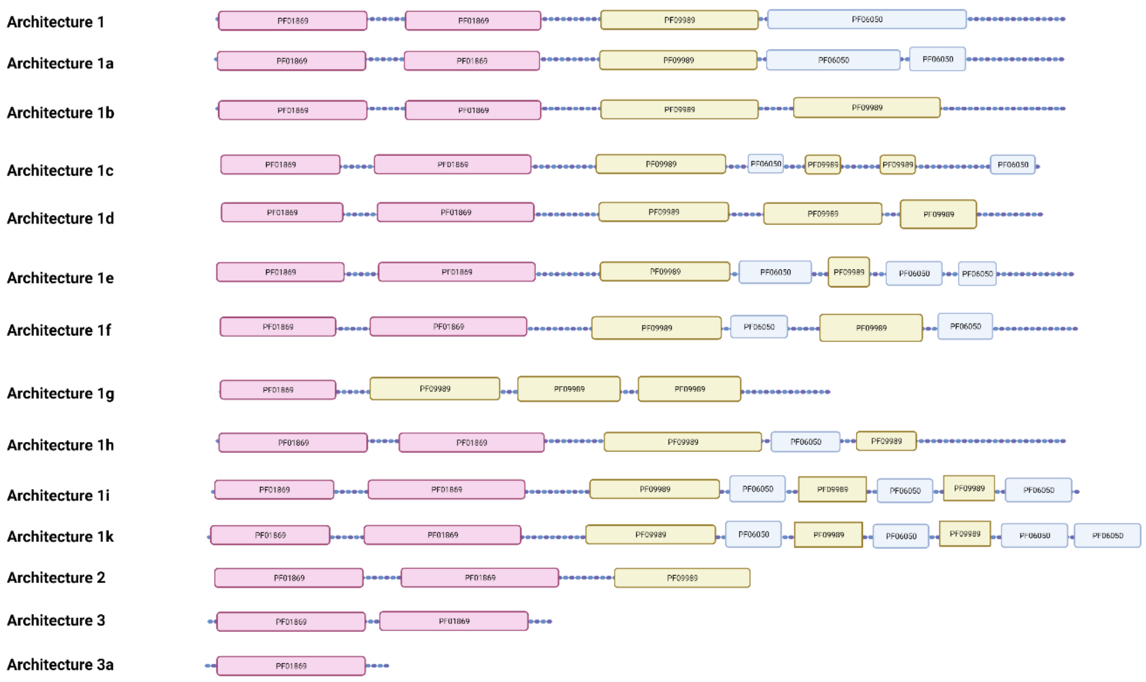

#### Supplementary Tables (provided as a compiled excel file)

#### Supplementary Files

**Supplementary File 1.** Maximum likelihood phylogenetic tree of the predicted homologs of the confirmed MSL synthase across the tree of life grouped at the phylum or superphylum level as shown with the colors on the branches and indicated in the legend. See Figure 5a for further details.

**Supplementary File 2.** Maximum likelihood phylogenetic tree of the predicted homologs of the confirmed ether bond-forming enzyme, GeR, across the tree of life grouped at the phylum or superphylum level as shown with the colors on the branches and indicated in the legend. See Figure 5b for further details.

**Supplementary File 3.** Negative sequence identities list for pBLAST searches of homologs to the selected upregulated radical SAM proteins from *T. ethanolicus*
