## Supplementary figures and images for "Disentangling the lipid divide: Identification of key enzymes for the biosynthesis of unusual Membrane-spanning and Ether lipids in Bacteria"

### Supplementary File 1

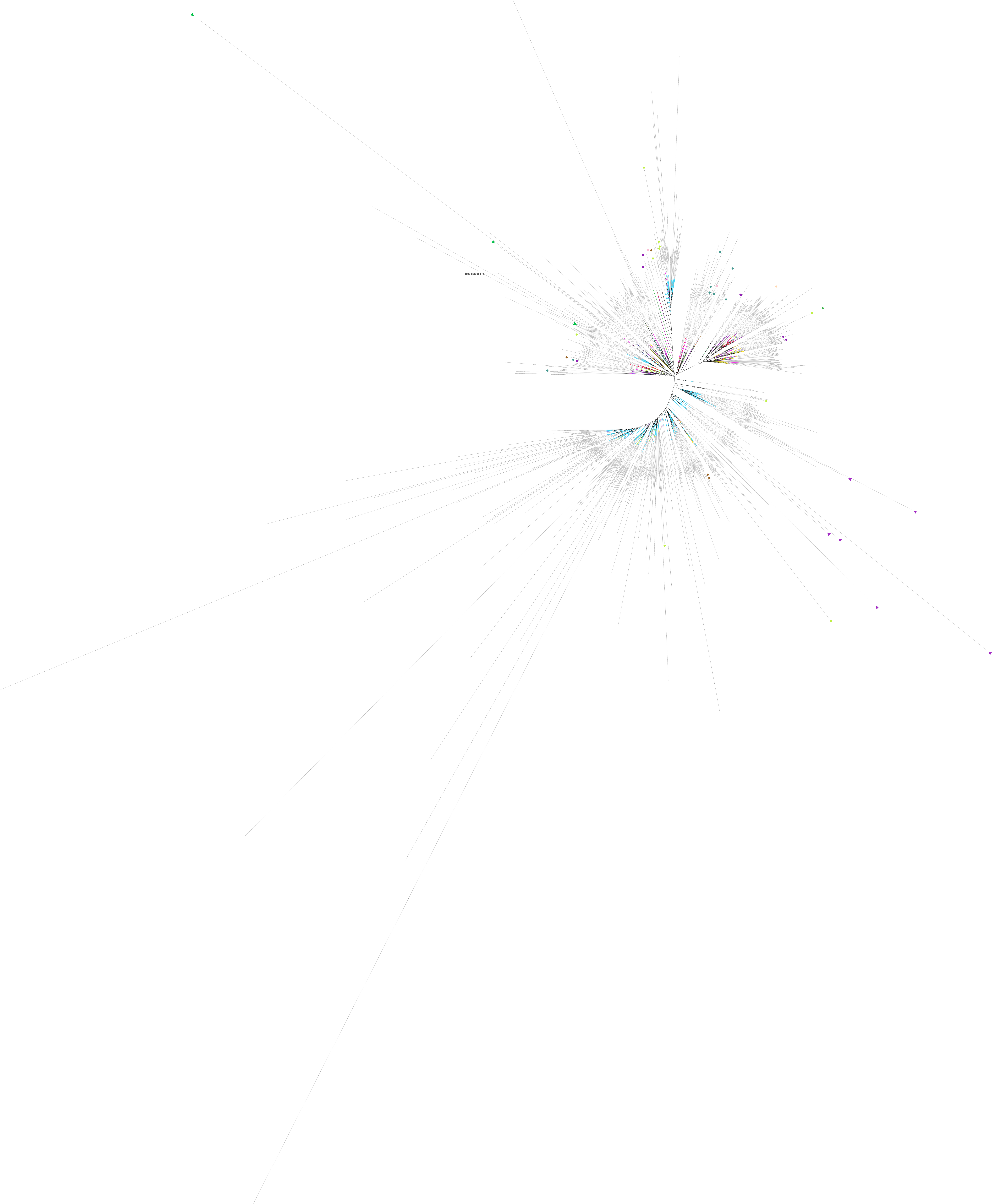

### Supplementary File 2

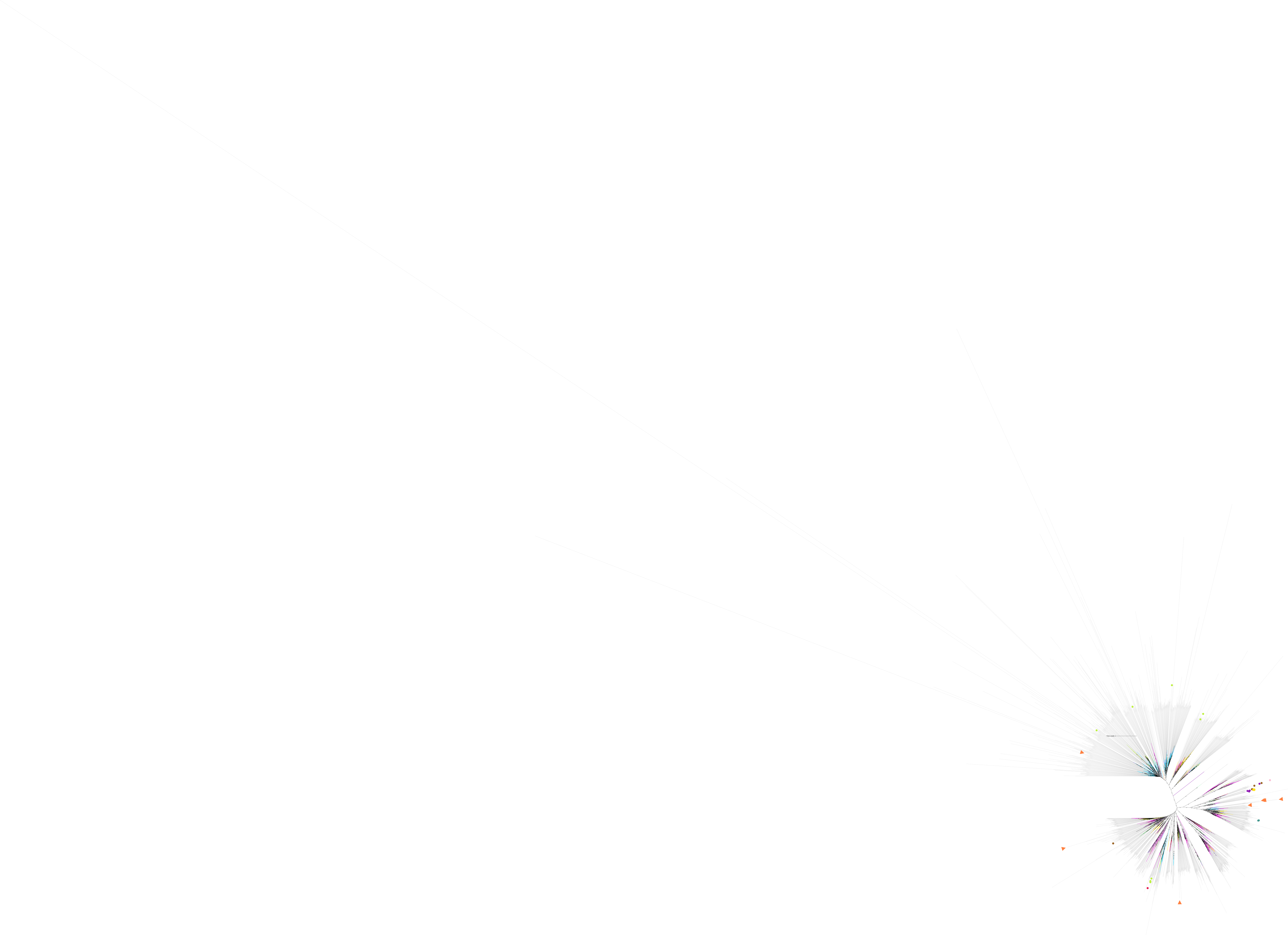
